# Modes of natural selection on maternal and zygotic gene expression in *Drosophila melanogaster* embryos

**DOI:** 10.64898/2026.05.29.728885

**Authors:** Thomas S. O’Leary, Brent L. Lockwood

## Abstract

Early embryonic development involves the coordination of gene expression from two distinct genomes, as mothers load eggs with gene products prior to the beginning of zygotic transcription. Because the maternal transcriptome is controlled by the mother’s regulatory genotype rather than that of the embryo, these two sequential developmental programs may experience distinct evolutionary constraints and selection pressures despite both existing within the embryo. To infer modes of selection on gene expression at maternal and zygotic developmental stages, we used single-embryo RNA-sequencing and conducted variance tests of selection on transcriptomes of parents and F_2_ segregants of tropical and temperate *Drosophila melanogaster* embryos. We found that approximately 10% of maternal transcripts and 5% of zygotic transcripts showed signals of selection. Genome-wide, directional selection was more common than stabilizing selection. However, among core early developmental genes and transcription factors, maternal transcription showed patterns of both stabilizing and directional selection, whereas zygotic transcription was predominantly under stabilizing selection. Many heat shock genes showed patterns of directional selection between tropical and temperate embryos, consistent with local adaptation. Additionally, directional selection in piRNA-pathway genes suggests a role for germline defense during embryogenesis. Overall, while our data support the canonical view that core developmental networks are constrained by stabilizing selection, the genome-wide prevalence of directional selection highlights a substantial and previously underappreciated contribution of diversifying selection to the evolution of early development.

**Article Summary:** The evolution of embryonic gene expression remains poorly characterized on microevolutionary timescales. We conducted genetic crosses between tropical and temperate *Drosophila melanogaster*, combining single-embryo RNA sequencing with variance tests of selection, to infer patterns of adaptive evolution in gene expression. We found pervasive signals of selection. While core developmental genes were largely under stabilizing selection, directional selection was the predominant mode of selection genome-wide. These findings challenge the paradigm that early embryogenesis is evolutionarily inflexible, suggesting that diversifying selection and local adaptation play key roles in the evolution of development on short evolutionary timescales.

## Introduction

The evolutionary constraints shaping early development remain poorly understood despite the central role of embryogenesis in organismal fitness (González-Forero, 2023; Kalinka and Tomancak, 2012). This knowledge gap may reflect the complexity of development itself, which involves interactions among numerous genetic factors (Davidson, 2006). Meanwhile, developing cells and tissues must meet strict functional requirements as they differentiate and organize (Ko and Martin, 2020).

Adding to this complexity, early development relies on the temporal coordination of two genetic components: the maternal genome and the zygotic genome (Vastenhouw et al., 2019). In all animals, genetic control of embryogenesis hands off from maternal factors to the zygotic genome during the maternal-to-zygotic transition (MZT) (Tadros and Lipshitz, 2009; Vastenhouw et al., 2019). In *Drosophila melanogaster*, the spatial gradients guiding early development comprise maternal mRNAs and proteins (*e*.*g*., *bicoid*) loaded by maternal nurse cells during oogenesis (Driever and Nüsslein-Volhard, 1988; Schüpbach and Wieschaus, 1986). These maternal products guide the earliest stages of embryonic development, establishing axial patterning and driving the initial waves of mitotic division (Edgar and O’Farrell, 1989; Foe and Alberts, 1983). As development proceeds, many maternal transcripts are degraded, and the zygotic genome becomes transcriptionally active during zygotic genome activation (ZGA). This activation drives the gene networks that specify primordial tissues and prepare the embryo for gastrulation (Berg et al., 2024; Crews, 2019; Gheisari et al., 2020; Martin, 2020; Öztürk-Çolak et al., 2018; Pollex et al., 2024).

This temporal progression may lead to shifting evolutionary constraints across embryogenesis, as developmental gene networks become increasingly interconnected (Davidson, 2006), while the passing of genetic control from maternal to zygotic inputs may change the specific gene targets available for natural selection (Atallah and Lott, 2018). Accordingly, comparative studies across diverse taxa—including insects (Liu et al., 2021), vertebrates (Irie and Kuratani, 2011), and plants (Quint et al., 2012)—show that morphological and molecular divergence between species varies across developmental time.

While extensive work has characterized phenotypic and molecular variation throughout development (Davidson, 2006; Raff, 1996), most studies have focused on divergence between species with comparatively little attention to variation within species (Coronado-Zamora et al., 2019; Feitzinger et al., 2022).

Consequently, modes of selection—stabilizing versus directional—have typically been inferred from relative rates of interspecific divergence rather than through explicit intraspecific tests of selection (Coronado-Zamora et al., 2019). Furthermore, the relative contributions of maternal versus zygotic gene products to natural genetic variation have only recently been investigated (Feitzinger et al., 2022; Omura and Lott, 2020). Thus, the extent to which selection differentially shapes maternal and zygotic transcriptional variation within a species remains unclear.

Direct tests of selection on gene expression are challenging because they require a baseline of neutral evolution against which to identify signals of directional and stabilizing selection (Price et al., 2022). A simple genetic crossing design solves this problem by leveraging random recombination in F_2_ progeny to generate a useful null distribution (Fraser, 2020). Under this framework, parental divergence smaller than the expression variance among F_2_ segregants indicates stabilizing selection. Conversely, parental divergence exceeding F_2_ variance is evidence of directional selection. This design allows the characterization of selection operating over relatively short evolutionary timescales, which is relevant to local adaptation.

In the present study, we used reciprocal crosses to generate F_2_ segregants from two natural isolates of *D. melanogaster*: one isofemale line from the tropical island of St. Kitts in the Caribbean Sea, and another from temperate North America in Vermont, USA. These two lines share genetic ancestry but have evolved in distinct environments for hundreds to thousands of generations (Haudry et al., 2020; Nunez et al., 2025; Nunez et al., 2026) (**Fig. S1, A and B**). We hand-staged embryos at maternal (Bownes’ stage 2) and zygotic (Bownes’ stage 5) expression timepoints and quantified transcript levels using single-embryo RNA-seq. Using a variance-based selection test, we inferred whether maternal and zygotic expression of each gene showed evidence of stabilizing or directional selection (**Fig. S1, C–F**).

We formulated three general predictions regarding these selection patterns:

First, we predicted that stabilizing selection would be more prevalent than directional selection, regardless of developmental stage. Gene expression is likely to be maintained near an evolutionary optimum over time (Bedford and Hartl, 2009; Rifkin et al., 2005). Indeed, small changes in transcript abundance (de Lachapelle and Bergmann, 2010; Reeves and Stathopoulos, 2009; Struhl et al., 1989) or timing (Dufourt et al., 2018; McDaniel et al., 2019) can severely disrupt embryogenesis. Accordingly, embryonic transcriptomes are generally conserved across *Drosophila* species (Atallah and Lott, 2018), which may be the result of stabilizing selection acting to preserve consistent developmental outcomes (Waddington, 1942). And, previous studies to quantify selection in early development only report evidence for stabilizing selection or purifying selection (Coronado-Zamora et al., 2019; Liu and Robinson-Rechavi, 2018).

Second, we predicted that maternally deposited transcripts would show more widespread signals of selection than zygotic transcripts. The “hourglass” model of developmental divergence (Duboule, 1994) is consistent with this prediction, as the earliest stages of development show a greater degree of phenotypic variation among species than intermediate stages (Raff, 1996). In fact, developmental transcriptomes across the *Drosophila* phylogeny follow this trend, exhibiting elevated levels of polymorphism and divergence at the earliest embryonic stage (Coronado-Zamora et al., 2019; Kalinka et al., 2010). This macroevolutionary pattern indicates that the maternal program is less evolutionarily constrained than the subsequent zygotic program. If a similar landscape of constraint shapes variation within *D. melanogaster*, the maternal transcriptome should harbor more standing variation for selection to act upon than the zygotic transcriptome. Additionally, because the maternal transcriptome is primarily controlled by *trans*-acting factors and chromatin state (Omura and Lott, 2020), selection on a small number of upstream regulators can shift expression phenotypes across downstream gene networks, which would inflate the proportion of genes with detectable signals of selection. In this manner, the realized effect of selection may be greater on maternal transcripts than zygotic transcripts simply due to network amplification.

Third, because our genetic lines were sourced from populations that evolved in contrasting tropical and temperate climates, maternal gene products may reflect adaptive differences in thermal tolerance (Lockwood et al., 2017; Lockwood et al., 2018). Stage 2 embryos have a reduced zygotic transcriptional response to heat stress (Graziosi et al., 1980; Mikucki et al., 2024); thus, maternally deposited transcripts may represent a critical aspect of the early embryo’s defense against thermal stress. Therefore, directional selection on thermally protective maternal transcripts is a plausible mechanism underlying population-level differences between St. Kitts and Vermont lines (Lockwood et al., 2017; Lockwood et al., 2018; Nunez et al., 2026).

Our results allow us to evaluate these predictions and provide a transcriptome-wide perspective on the role of selection in shaping gene expression during early embryonic development in *D. melanogaster*.

## Materials and methods

### Fly care

For several generations prior to experimentation, we raised the two parental fly lines, St. Kitts (SK) and Vermont (VT), on the same standard yeast, cornmeal, and molasses food. The SK parental line was collected in Monkey Hill, St. Kitts and sourced from the *Drosophila* Species Stock Center at the University of California (stock no. 14021-0231.34). The VT parental line was collected in East Calais, Vermont, USA (Cooper et al., 2014; Lockwood et al., 2018). Both parental lines were established with a single female founder and isogenized through several generations of full-sibling mating. The stock vials were maintained at a standard density of approximately 50 to 100 flies per vial (95 mm × 25 mm, Genesee Scientific) and were kept in an incubator (DR-36VL, Percival Scientific Inc.) at 25°C and 55% relative humidity on a 12hr:12hr light:dark cycle.

### Fly crosses to create F_2_ segregants

We performed reciprocal crosses of the parental lines (☿_SK_ × ♂_VT_ and ☿_VT_ × ♂_SK_) to produce F_1_ hybrids (**Fig. S1, A and B**). Reciprocal F_0_ crosses ensured that downstream cohorts contained both St. Kitts and Vermont mitochondrial haplotypes to balance the maternal-line cytoplasmic background across F_2_ segregants. After the F_1_ hybrid flies eclosed, the adults were split into two cohorts. One cohort was used immediately to collect F_2_ segregant embryos for the variance test on the zygotic stage of expression (Bownes’ stage 5). The other cohort was interbred (☿_F1_ × ♂_F1_) in separate food vials to raise F_2_ segregant mothers that were used to lay eggs in collection cages (☿_F2_ × ♂_F2_) for the variance test on a maternal stage of expression in embryos (Bownes’ stage 2).

### Embryo collection and staging

For both the F_2_ segregants and the parentals, egg collection cages (Genesee Scientific) were established with approximately 100 mating pairs with grape juice agar plates (60 mm × 15 mm) with yeast paste. The flies acclimated to their cages for two days prior to experimental egg collection, and plates were refreshed with fresh grape juice agar plates with yeast paste every 24 hours. Immediately before collection, for two successive hours, we supplied the fly cages with new egg-laying substrate (*i*.*e*., fresh grape juice agar plate with yeast paste) to purge eggs of unknown age held by mothers. On a fresh grape juice agar plate with yeast, we collected embryos for 15 minutes. Agar plates with embryos for the zygotic expression samples were left in the incubator to develop for 2 hours and 45 minutes, but the embryos for maternal expression were processed immediately. The embryos were washed (0.7% (*w/v*) NaCl, 0.05% Triton X-100) and then rinsed in R.O. H_2_O. The embryos were submerged in 50% bleach for one minute for dechorionation, and then immediately rinsed with R.O. H_2_O. Each individual embryo was transferred with a paintbrush to a slide and checked for the appropriate embryonic developmental stage using a Leica M80 Stereo Zoom Microscope. Maternal expression embryos (Bownes’ stage 2) were defined as embryos that have not yet formed pole cells. Zygotic expression embryos (Bownes’ stage 5) were defined as embryos that had formed pole cells and undergone cellularization of the blastoderm but had not yet gastrulated. Once the developmental stage was confirmed, we added a drop of 0.5 µL TRIzol (Tri-reagent, Sigma Life Science, St Louis, MO, USA) on top of the embryo. We then punctured the embryos with a clean 23-gauge needle (Kendall, Monoject) and allowed the embryo to dissolve in the TRIzol for 5 minutes. The drop of TRIzol with the dissolved embryo was transferred to a 1.5 mL microcentrifuge tube with 1 mL of TRIzol. The spot on the slide was rinsed with 1 mL TRIzol to pick up any remaining fluid or debris and added to the sample tube. The samples were immediately flash frozen in liquid nitrogen and stored at –80°C.

### RNA extraction

Sample tubes with 1 mL TRIzol were gently thawed on top of ice. We then added 200 µL chloroform to each tube, mixed by shaking, and incubated the samples on ice for 3 minutes. We spun the samples at 12,000 RCF for 15 mins at 4°C (Sorvall ST89, Thermo Scientific). The top 400 µL of the aqueous phase containing RNA was transferred to a fresh 1.5 mL tube. To aid with precipitation, we added 8 µL of 20 µg/µL glycogen (Thermo Scientific), 500 µL isopropanol (Thermo Scientific), and incubated the sample on ice for 10 minutes. We then centrifuged the samples at 12,000 RCF for 15 mins at 4°C and discarded the supernatant. We washed the pellet in 1 mL 75% ethanol, and centrifuged the sample at 7,500 RCF for 5 mins at 4°C. The supernatant was discarded, and we air-dried the pellet. The sample was resuspended in 20 µL of nuclease-free H_2_O and incubated for 10 minutes at 60°C. Samples were checked for quality control with a NanoDrop^™^ 2000 (Thermo Fisher) and 2100 BioAnalyzer (Agilent) prior to shipping them for sequencing. Two samples (one Bownes’ stage 5 SK sample, and one Bownes’ stage 2 F_2_ segregant sample) were discarded after failing to meet the quality control criteria for Novogene after shipping and handling. One sample (the Bownes’ stage 2 F_2_ segregant) showed signs of RNA degradation on the BioAnalyzer trace, and the other sample (Bownes’ stage 5 SK) had an abnormal trace pattern. This left 15 Bownes’ stage 2 samples (3 of each parental line and 9 F_2_ segregants) and 21 Bownes’ stage 5 samples (5 SK, 6 VT, and 10 F_2_ segregants). We chose to include more parental Bownes’ stage 5 embryos because we wanted to increase the sample of both male and female embryos, as this is a stage in embryogenesis where sex-specific expression has begun (discussed in more detail below).

### Library preparation & RNA-sequencing

Library preparation and RNA-sequencing was completed through Novogene. Briefly, approximately 5 to 25 ng of total RNA was used for library preparation using the SMART-Seq® v4 Ultra® Low Input RNA Kit (Takara Bio USA, Inc.) according to manufacturer’s specifications. Libraries were sequenced on an Illumina NovaSeq X Plus Series for 150 bp paired-end targeting 6 G per sample of total depth.

### Quality control, mapping, and data normalization

Raw fastq files were analyzed with FastQC (version 0.12.1) and the output was aggregated across all samples with MultiQC (version 1.28). Reads were quasi-mapped with salmon (version 1.10.0; (Patro et al., 2017) to the reference transcriptome from Ensembl (BDGP6 assembly, annotation release r6.46; file Drosophila_melanogaster.BDGP6.46.cdna.all.fa). The count matrices for each sample were uploaded to R (version 4.5.0; (R Core Team, 2025). The data were normalized with variance-stabilizing transformation (VST) from DESeq2 (version 1.48.1; (Love et al., 2014). Only genes with reads in all samples of the same stage were used for the variance test for selection.

### Anomalous Bownes’ stage 5 samples

Principal component analysis revealed 5 samples that did not cluster with the other zygotic expression samples (Sample IDs: SZ8, SZ9, VZ1, VZ4, F2Z13; two SK, two VT, and one F_2_ segregant), we have termed these five samples anomalous. Because zygotic expression is rapidly ramping up at this stage (Vastenhouw et al., 2019), it is possible that small differences in the developmental stage of these embryos led to large differences in the composition of their transcriptomes. For the variance test, because we needed to focus on variation that may be due to selection and exclude variation due to differences in developmental timing, for the main text, we have chosen to focus on the 16 Bownes’ stage 5 embryos (3 SK, 4 VT, and 9 F_2_ segregant samples) that cluster together in PCA space and show patterns of sex-specific expression (another indication of the developmental timing disconnect with these 5 anomalous samples – see more information in the section below). However, we are including downstream analyses for all 21 Bownes’ stage 5 samples (5 SK, 6 VT, and 10 F_2_ segregant samples) in the supplement.

### Sex-specific expression in Bownes’ stage 5 zygotic expression embryos

We determined the sex of the Bownes’ stage 5 embryos using the expression of two genes: *msl-2* and *Sxl*. Male (XY) embryos have high expression of *msl-2* and low expression of *Sxl* and female (XX) embryos have the inverse pattern (Pérez-Mojica et al., 2023). The five anomalous zygotic expression samples did not have high enough expression of these two genes and were excluded from this analysis. To determine which genes displayed sex-specific expression patterns, we ran DESeq2 between males and females. Genes that displayed sex-specific expression (adj. p < 0.05) were excluded from the variance test for selection in the Bownes’ stage 5 samples.

### Nonparametric variance test for selection

For genes that violated the assumption of a normal distribution in the F_2_ segregants (Shapiro-Wilk; p-value < 0.05), we performed a nonparametric version of the variance test following (Kern et al., 2021), using a rank-based dispersion comparison. We first quantified the spread of the F_2_ segregants as the absolute deviation of each F_2_ sample from the mean of the F_2_ segregants. To estimate the parental dispersion, we formed all non-repeating SK–VT pairs, calculated each pair’s mean, and took absolute deviations from its pair mean. We then applied a Kruskal–Wallis rank-sum test to compare the two sets of dispersion values. To create a null distribution, we pooled all parental and F_2_ segregant samples and permuted the group labels 20,000 times and recomputed the Kruskal–Wallis statistic. Stabilizing selection p-values were calculated as the proportion of permuted test statistics less than or equal to the observed test statistic. Directional selection p-values were calculated as the proportion of permuted test statistics greater than or equal to the observed test statistic. The p-values for the non-parametric test were pooled with the p-values from the parametric version, and the false discovery rate was corrected for using the Benjamini–Hochberg procedure.

To compare the effect-size of selection across developmental stages, we calculated a median dispersion ratio (MDR) for each gene by dividing the median parental dispersion by the median F_2_ dispersion. In this way, MDR values much greater than one indicate a large directional signal and values close to zero indicate strong stabilizing selection. Parental log_2_(fold-differences) were calculated as a ratio of the mean Vermont expression to St. Kitts, so positive values indicate higher Vermont expression and negative values indicate higher St. Kitts expression.

### Estimating non-additive genetic effects

We define non-additivity as a significant deviation of the F_2_ mean expression from the midparent mean expectation under additive inheritance. Biologically, this can arise from several distinct mechanisms, including epistatic gene interactions and dominance. We estimated non-additive effects (Δ) using **Eq. 1**, where μ_F2_ represents the mean expression value for the F_2_ segregants and μ_VT_ and μ_SK_ represent the mean expression of the parental lines (Vermont and St. Kitts respectively). Positive Δ values indicate that the F_2_ mean exceeds the midparent expectation, and negative Δ values indicate F_2_ mean is lower than the additive expectation (Fraser, 2020). Following (Kern et al., 2021) to create a t-statistic, we divided Δ by the square root of the standard error of the mean (**Eq. 2 & Eq. 3**). Next, we permuted the group labels 20,000 times to create permuted t-statistics and calculated the p-value as the fraction of permuted t-values that were greater in magnitude than the absolute value of the observed t-statistic. P-values were adjusted to correct for multiple comparisons using the Benjamini–Hochberg procedure.

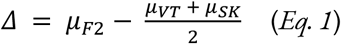

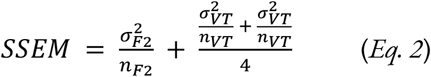

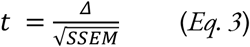

Additionally, for directionally selected genes displaying non-additivity, we quantified the degree of parental bias by calculating a parental bias index (PBI, **Eq. 4**). This index is centered and scaled, so that 0 indicates the midparent expectation, negative values indicate bias towards St. Kitts, positive values indicate bias towards Vermont, and any values greater than the absolute value of 1 indicate F_2_ mean outside of the parental range.

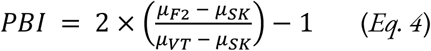

### Gene ontology (GO) over-representation analysis

We conducted an over-representation analysis (ORA) using WebGestalt (Elizarraras et al., 2024) on the genes with a significant signal for selection for maternally deposited and zygotically expressed genes. For directionally selected genes, we also separately tested genes that had been directionally selected for higher expression in the Vermont or St. Kitts embryos. We separately tested against the non-redundant Biological Process, Cellular Component, and Molecular Function databases. The ORA was conducted using many of the default parameters from WebGestalt unless otherwise specified. All GO terms with 5 to 2,000 elements were included in the analysis and GO terms were marked as significantly over-represented when they passed false discovery rate correction (Benjamini-Hochberg adj. p < 0.1).

### Targeted gene-set enrichment analysis

In addition to the GO over-representation analysis, we tested whether predefined, curated gene sets were over- or under-represented among genes under selection. We focused on three sets: (i) canonical embryogenesis genes relevant to Bownes’ stage 2 and 5 embryos, (ii) transcription factors (TFs), and (iii) heat shock proteins (HSPs). For each developmental stage separately and for each gene set, we formed a 2 × 3 table comparing the membership in the gene set and selection class (stabilizing, neutral evolution, and directional) versus the background of all other genes tested to which we applied a two-sided Fisher’s exact test. If the test was significant, we then applied a two-sided Fisher’s exact test to compare directional or stabilizing selection separately to neutral evolution in a 2 × 2 table.

TFs were taken from the FlyBase gene group “Transcription Factors” (FBgg0000745, date last reviewed: June 1, 2017) containing 628 genes; 456 TFs were tested, with 330 in the maternal and 427 in the zygotic. HSPs were taken from the FlyBase gene group “Heat Shock Proteins” (FBgg0000501, date last reviewed: April 21, 2016) containing 86 genes; a total of 67 HSP genes tested for selection, with 62 in the maternal 64 in the zygotic. Canonical embryogenesis genes were curated from The Interactive Fly (Brody, 1999). For the maternal gene set (Bownes’ stage 2), we gathered genes from the Anterior group, Posterior group and the formation of the germ plasm, Terminal group, and Dorsal group (Brody, 2024a) for a total of 68 genes; 61 of these genes cleared expression filters genes were tested for selection in maternal expression. For the zygotic gene set, we gathered genes from the Gap genes, Pair rule genes, and Dorsal-Ventral Patterning Genes and Dpp Signaling (Brody, 2024b) for a total of 99 genes, 82 of which were among the genes tested for selection.

## Results

### Sequencing quality control

In total, we generated 1.06 billion 150 bp paired-end reads across 36 samples (mean = 29.36 million reads). On average, 80.6% of reads mapped, yielding 23.67 million mapped reads per sample (**Fig. S2A**). These reads mapped to an average of 8,990 genes per sample (**Fig. S2B**). Gene detection differed significantly between stages (Welch’s two-sample t-test; t = –8.23, p = 2.7 × 10^−8^): zygotic (Bownes’ stage 5; mean = 9,683) samples had 20% more genes than maternal samples (Bownes’ stage 2; mean = 8,020). In this manuscript, we will refer to Bownes’ stage 2 as maternal expression and Bownes’ stage 5 as zygotic expression, because these stages align with before and after the maternal-to-zygotic transition (Tadros and Lipshitz, 2009; Vastenhouw et al., 2019). Transcripts present at stage 2 reflect maternal deposition, whereas transcripts present at stage 5 likely represent some combination of newly activated zygotic transcription and maternally deposited transcripts that have not yet been degraded (Atallah and Lott, 2018). In total, 10,363 genes were detected in the maternal samples, and 12,869 genes were detected in the zygotic expression samples. Although we captured many low-expression genes, the majority had more than 100 counts across all samples (**Fig. S2, C and D**). After filtering for only genes included in the downstream variance test, the median per-gene read count was 1,990.5 for maternal and 869 for zygotic samples (**Fig. S2, E and F**).

### Principal component analysis and identification of five anomalous zygotic samples

After variance-stabilizing transformation (Love et al., 2014), principal component analysis (PCA) showed that the 15 maternal samples were clustered tightly, whereas the 21 zygotic samples were much more dispersed (**Fig. S3A**). PC1 captured 38% of the variance, and largely separated the stages, with all 15 maternal expression samples with negative PC1 scores, and 16 out of the 21 zygotic samples had positive PC1 scores (**Fig. S3B**). Despite careful egg-collection timing and staging, we suspect that small differences in developmental timing resulted in anomalous expression patterns in five embryos (Vastenhouw et al., 2019). Accordingly, we conducted all zygotic analyses both with and without these five samples (see methods for additional details). Unless otherwise noted, zygotic results in this manuscript refer to the filtered set (n = 16) without the five anomalous samples and the analyses with the full set of zygotic embryos (n = 21) are included in the supplement. Excluding the five anomalous samples, PCA clustering of all samples remained largely unchanged (**Fig. S3C**), with PC1 now explaining 44% of the variance (**Fig. S3D**).

### Stage-specific principal component analysis

To examine within-stage variation, we conducted a PCA separately on the maternal and zygotic samples. In the maternal PCA, PC1 captured 23% of the variance and clearly separated the Vermont embryos from the St. Kitts and F_2_ segregants which overlapped (**Fig. S4, A and B**). For the zygotic expression including the five anomalous samples, PC1 explained 31% of the variation and separated out the anomalous samples from the other 16 (**Fig. S4, C and D**). After excluding the anomalous samples, the Vermont, St. Kitts, and F_2_ segregant embryos predominantly overlapped on the first two PCs (**Fig. S4E**), and the variance was more evenly distributed across the first several PCs (**Fig. S4F**).

### Sex-specific expression of zygotic embryos

To control for sex-specific zygotic expression, we assigned sex to each zygotic sample using *msl-2* and *Sxl* expression following (Pérez-Mojica et al., 2023). Of the 21 samples, 16 could be sexed, but the five anomalous samples lacked sufficient expression levels for confident assignment (**Fig. S5A**), consistent with the PCA results presented above. We identified 7 females and 9 males and used these groups to define a list of sex-specific genes to exclude from the zygotic expression variance test. Differential expression analysis (DESeq2 Wald test; adj. p < 0.05) found 290 genes with sex-specific expression patterns (**Fig. S5B)**. There were 211 genes with higher expression in female embryos and 79 genes higher in males, with *Sxl* and *msl-2* genes showing the strongest difference. Most sex-specific genes (84.1%) mapped to the sex chromosomes with 215 genes (74.1%) on the X and 29 genes on the Y (10.0%) despite the small number of genes it encodes (**Fig. S5C**). We did not have enough within-sex replication to test for selection on expression levels within male and female embryos separately, so all sex-biased genes were removed from downstream analyses. Therefore, our estimates of selection may underestimate the true level selection on gene expression, as sex-biased genes are known to evolve rapidly (Ellegren and Parsch, 2007; Grath and Parsch, 2016).

### Genes included in the variance test

To focus our selection analysis, we retained only genes with mapped reads in every sample, yielding 6,952 maternal and 7,869 zygotic genes (all zygotic analyses refer to the filtered set unless noted, see above). Of the 8,330 total genes tested, the vast majority (6,491; 77.9%) were common to both stages, while 1,378 (16.5%) were unique to the zygotic samples and 461 (5.5%) were unique to the maternal samples (**Fig. S6A**). For the purposes of this study when we refer to detected maternally deposited genes or zygotic genes, we are referring to the set of genes that passed the above expression filters. These genes may be shared (*i*.*e*., present at both stages) or stage-specific (*e*.*g*., maternal-only or zygotic-only), meaning the gene is detected at one stage but absent from the other.

### Detecting selection on gene expression

Of the 6,952 maternally deposited genes, 193 (2.8%) showed stabilizing selection and 517 (7.4%) showed significant directional selection — summing to 10.2% of maternally deposited transcripts. Among the 7,869 zygotic genes, 60 (0.8%) were under stabilizing selection and 368 (4.7%) were under directional selection — totaling 5.4% of zygotic transcripts (**Fig. 1A**). Maternal and zygotic genes had different distributions of selection hits (Fisher’s exact test, p = 1.0 × 10^−6^), with maternally deposited genes having disproportionately more stabilizing selection (Fisher’s exact test, odds ratio = 3.83 [95% CI 2.9–5.2], p = 1.2 × 10^−22^) and directional selection (Fisher’s exact test, odds ratio = 1.67 [95% CI 1.5–1.9], p = 1.7 × 10^−13^) than zygotically expressed genes. Despite the large number of shared genes, there were only a few genes that were under selection in both stages. Only one (<1%) stabilizing selection gene (*ND-MNLL*) was shared between the two stages (**Fig. S6B**), and only 75 directional selection genes (9.3%) were shared between the two stages (**Fig. S6C**).

**Figure 1.**
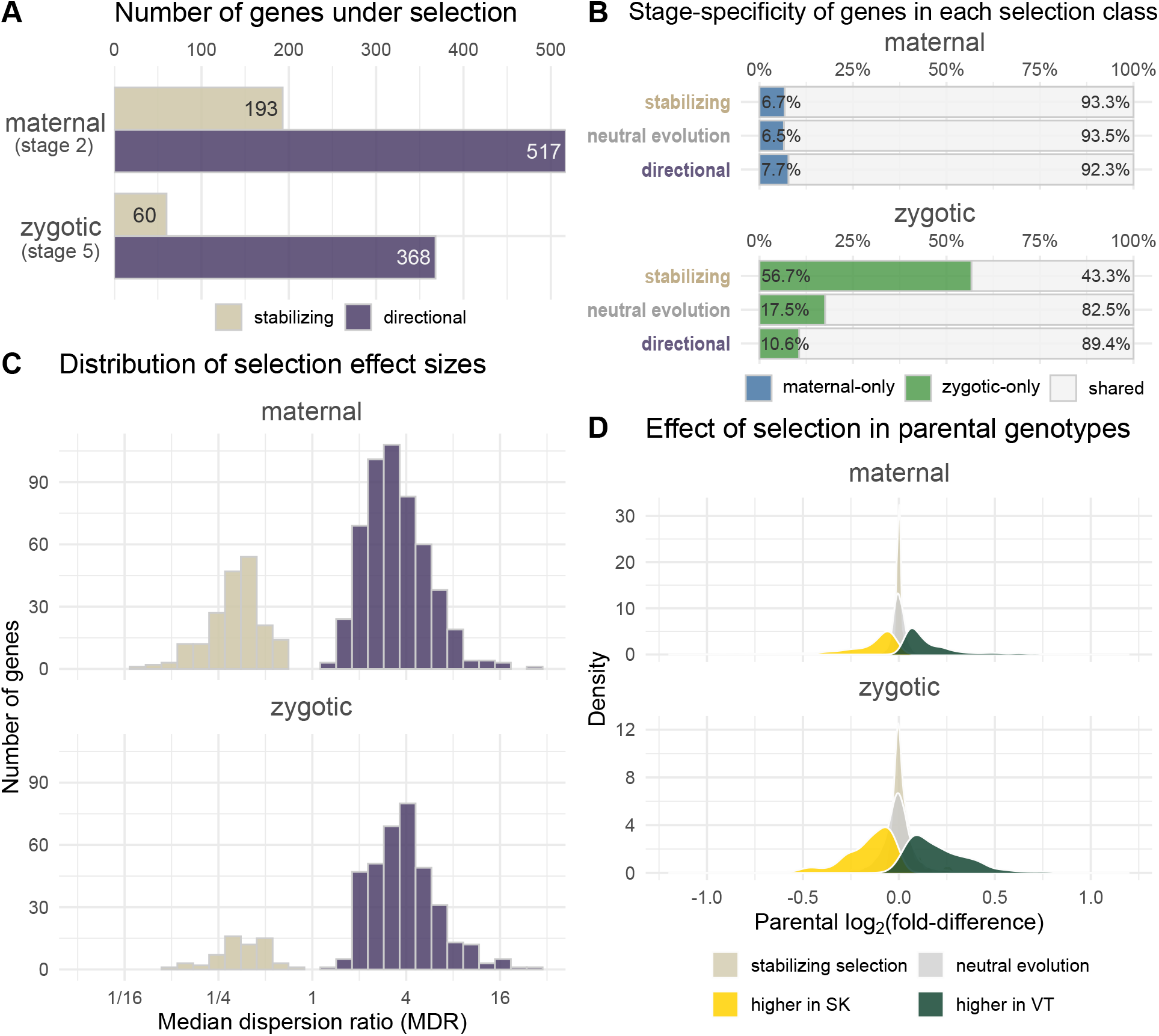
Selection is more prevalent among maternally deposited transcripts, while zygotic-only genes are enriched for stabilizing selection, and directional selection is balanced between St. Kitts and Vermont. (**A**) Bar plot showing counts of genes with significant stabilizing or directional selection in maternal (stage 2) and zygotic (stage 5) expression. (**B**) Bar plot showing the proportion of stage-specific genes in each selection class in maternal and zygotic expression. (**C**) Histogram of median dispersion ratio (MDR) for genes with significant selection signal; MDR > 1 indicates higher parental variance (directional) and MDR < 1 indicates lower parental variance (stabilizing). (**D**) Density plot showing the distribution parental log_2_(fold-difference) of directionally selected genes colored by genotype with higher parental expression: St. Kitts (yellow), Vermont (green), and neutral evolution background genes (grey).

To examine whether stage-specific transcripts were disproportionately represented among selection hits, we narrowed our analysis to genes that were detected in only one stage and compared those selection rates to those in the genes present in both stages (shared genes). Maternal-only genes (n = 461) were distributed across selection classes at similar rates relative to shared genes (Fisher’s exact test, p = 0.538). In contrast, zygotic-only genes (n = 1,378) were disproportionately represented in the selection hits relative to shared genes (**Fig. 1B**; Fisher’s exact test, p = 1.0 × 10^−6^). Zygotic-only genes comprised 56.7% of stabilizing selection hits (34 / 60) compared with only 17.5% of neutrally evolving genes (Fisher’s exact test, odds ratio = 6.15 [95% CI 3.6–10.7], p = 1.4 × 10^−11^). Additionally, zygotic-only genes were under-represented in the directional selection hits (39 / 368) at 10.6% (Fisher’s exact test, odds ratio = 0.56 [95% CI 0.39–0.78], p = 3.8 × 10^−4^).

To compare the effect size of selection across developmental stages, we calculated a “median dispersion ratio” (MDR) for each gene by dividing the median pairwise replicate-based parental dispersion by the median F_2_ dispersion. We designed this metric to mimic the v statistic in Fraser (2020) while reflecting our non-parametric framework (Kern et al., 2021); the interpretation is unchanged: parental dispersion greater than the F_2_ dispersion (MDR > 1) is indicative of directional selection, while parental dispersion smaller than F_2_ dispersion (MDR < 1) is indicative of stabilizing selection. Although the maternal and zygotic stages have similar distributions of selection strength (MDR) among significant genes (**Fig. 1C**), gene-by-gene MDR values correlate weakly between stages (Pearson’s r = 0.12, r^2^ = 0.014, p = 2.8 × 10^−22^, n = 6,491; **Fig. S6D**). This result indicates that while the observed magnitude of selection on individual genes is comparable across the stages, the identity of the genes under selection is largely stage-specific, consistent with the small overlap of selection hits between stages **(Fig. S6, B and C**). Directional selection did not favor either parental line; genes with higher mean expression in St. Kitts and Vermont occurred in comparable numbers in both stages (**Fig. 1D**). In the maternally deposited directionally selected genes (n = 517), 50.7% (262) had higher expression in St. Kitts and 49.3% (255) had higher expression in Vermont; the zygotic set (n = 368) was similarly balanced — 51.1% (188) higher in St. Kitts versus 48.9% (180) higher in Vermont. Note that the variance test cannot distinguish selection for higher expression in one parental line from selection for lower expression in the other.

### Impact of the anomalous samples on our selection results

To assess whether the five anomalous samples, representing potentially mis-staged embryos, drove spurious results, we ran selection tests on zygotic expression both with and without them. Excluding these samples increased the number of genes meeting our all-sample expression criterion from 6,576 to 7,869. As a result, 29 of the 60 stabilizing-selection genes and 37 of the 368 directional-selection genes were detected only after removing the outliers (**Fig. S7, A and B**). Importantly, despite these differences in which genes passed the expression filter, estimates of selection strength were well correlated between the two analyses (R^2^ = 0.58; **Fig. S7C**).

### Over-represented gene ontology terms in selection genes

Among genes under stabilizing selection, there were 14 over-represented GO terms in the maternal genes and one in the zygotic genes (**Fig. S8A**). The GO term *positive regulation of RNA biosynthetic process* (GO:1902680) was over-represented in both stages, comprising 18 and 10 genes respectively; however, none of those genes under stabilizing selection were shared between the two stages. For genes directionally selected towards higher expression in Vermont embryos, four biological process GO terms were over-represented (three maternal, one zygotic), and both stages shared *organic acid metabolic process* (GO:0006082) with four shared directionally selected genes, *slgA, CG4594, CG17486*, and *CG18586* (**Fig. S8B**). No GO terms were over-represented among the genes directionally selected for higher St. Kitts expression (adj. p < 0.1). In the cellular component category, only *germ cell nucleus* (GO:0043073) was enriched with four genes (*piwi, osk, rhi*, and *vas*) displaying directional selection towards higher expression in the Vermont embryos (**Fig. 2A**).

**Figure 2.**
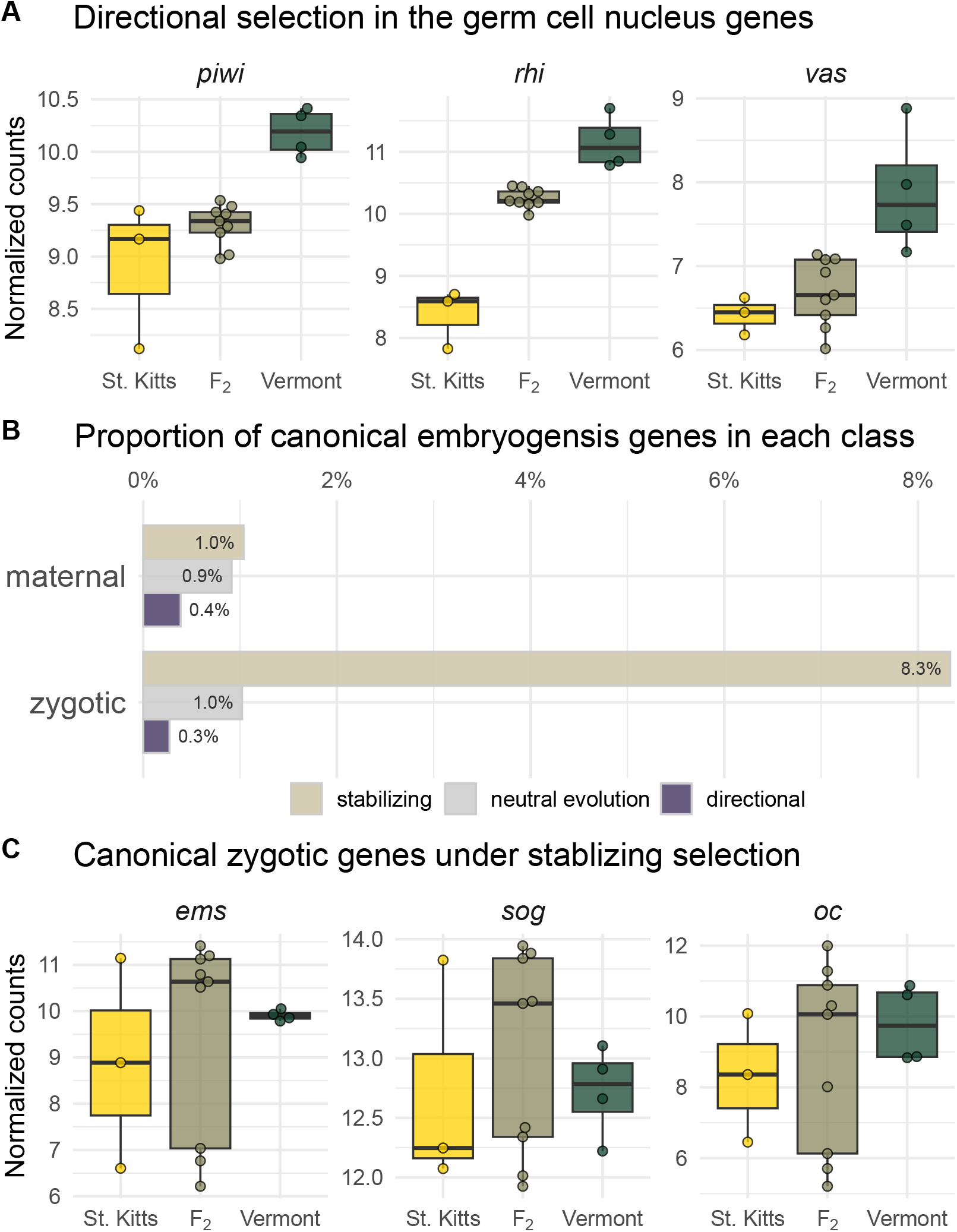
Germ cell nucleus genes show directional selection towards higher zygotic expression in Vermont embryos, whereas canonical zygotic embryogenesis genes are enriched for stabilizing selection. (**A**) Normalized expression of three germ cell nucleus genes (*piwi, rhi*, and *vas*) directionally selected for higher expression in Vermont embryos. Normalized expression values for individual embryos (points) overlaid on top of box plots (boxes represent the interquartile range; middle bar indicates the median; whiskers extend to 1.5×IQR). (**B**) Proportion of canonical embryogenesis genes among maternal and zygotic genes in each selection class. (**C**) Normalized expression of three canonical embryogenesis genes (*ems, oc*, and *sog*) active at Bownes’ stage 5 and under stabilizing selection.

### Selection on canonical genes involved in embryogenesis

To test whether the selection hits were enriched among the canonical genes known to be involved in development, we curated two lists of genes known to be involved in embryogenesis at Bownes’ stage 2 and 5 (see Methods). Canonical maternal genes (Bownes’ stage 2; n = 61) were present at similar rates among the selection classes compared with the background set of non-canonical development genes (**Fig. 2B**; Fisher’s exact test, p = 0.48); but the canonical zygotic genes (Bownes’ stage 5; n = 82) were disproportionately distributed in the selection hits compared to the background set (Fisher’s exact test, p = 2.6 × 10^−5^). Canonical genes comprised 8.3% of stabilizing selection hits (5 / 60) compared to only 1.0% of neutrally evolving genes (Fisher’s exact test, odds ratio = 10.1 [95% CI 3.5–23.8], p = 5.6 × 10^−5^). Although present at a lower rate of 0.3%, canonical zygotic genes were not statistically under-represented among the directional selection hits compared with neutrally evolving genes (Fisher’s exact test, odds ratio = 0.34 [95% CI 0.0403–1.25], p = 0.131). Among the canonical zygotic genes, 4 of the 5 genes selected for stabilizing expression are transcription factors involved in development (*cad, ems, oc*, and *cnc*), and the other gene (*sog*) is a dorsal-ventral patterning gene in the Dpp/BMP signaling pathway (**Fig. 2C**).

### Selection on the expression of transcription factors

We next examined selection levels across annotated transcription factors (TFs), due to their importance in regulating expression levels, a function of great importance during early embryogenesis and zygotic genome activation. Maternally expressed TFs (n = 330) were not enriched in the selection hits, comprising 6.2% of stabilizing hits (12 / 193) and 3.7% of directional hits (19 / 517) compared to 4.8% of neutrally evolving genes (299 / 5943; Fisher’s exact test, p = 0.30, **Fig. 3A**). However, zygotically expressed TFs (n = 427) were disproportionately distributed among the selection hits (Fisher’s exact test, p = 6.0 × 10^−6^), representing 20% of the stabilizing selection hits (12 / 60; Fisher’s exact test, odds ratio = 4.33 [95% CI 2.08–8.36], p = 9.3 × 10^−5^), but just 2.4% of the directionally selected genes (Fisher’s exact test, odds ratio = 0.43 [95% CI 0.20–0.84], p < 0.01), compared to 5.5% TFs among the neutrally evolving genes (5.5%; 406 / 7,441). TFs under stabilizing selection included core genes for early axis formation and segmental patterning such as *cad, cas*, and *Lim1* (**Fig. 3B**). Directionally selected TFs have evidence for a variety of functions including dorsal-ventral patterning and gastrulation via Dpp signaling (*fuss*) and mitochondrial synthesis and degradation (*Paris*) (**Fig. 3C**).

**Figure 3.**
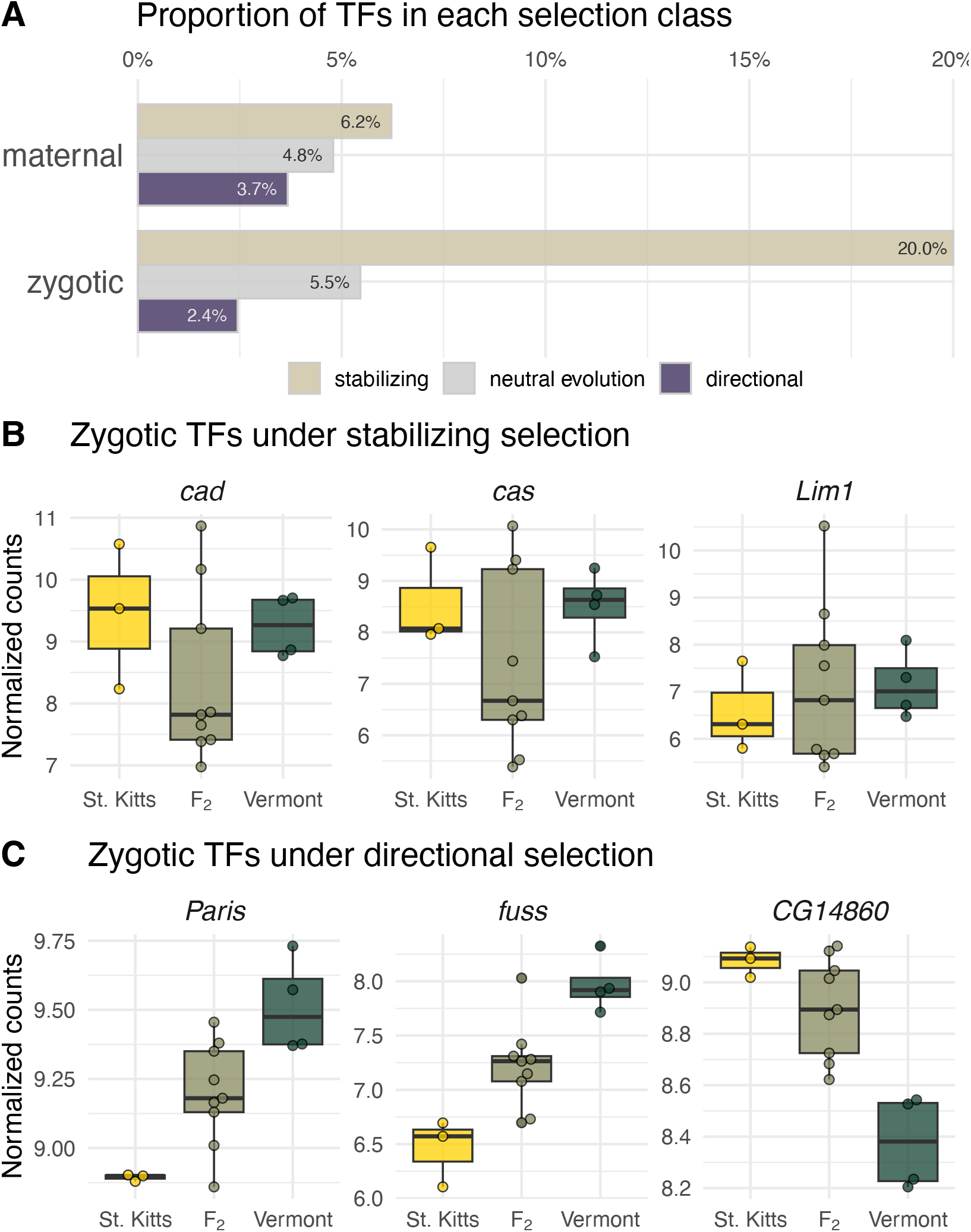
In zygotic expression, transcription factors are over-represented among stabilizing-selection genes and under-represented in the directional selection genes. (**A**) Proportion of TFs among maternal and zygotic genes in each selection class. (**B**) Normalized expression of three zygotic TFs, *cad, cas*, and *Lim1*, under stabilizing selection. (**C**) Normalized expression of three representative zygotic TFs, *CG14860, fuss*, and *Paris*, under directional selection. Normalized expression values for individual embryos (points) overlaid on top of box plots (boxes represent the interquartile range; middle bar indicates the median; whiskers extend to 1.5×IQR).

### Selection on heat shock proteins

We also chose to narrow our focus to the selection on heat shock protein expression, due to the evidence that increased maternal loading of heat shock proteins can affect heat tolerance in early embryos (Lockwood et al., 2017), and the observation that temperate and tropical early embryos differ in acute heat tolerance, a possible signature of thermal selection (Lockwood et al., 2018). Note that because our experiment was performed at 25°C and the variance test required non-zero counts in every sample, many heat-inducible HSPs (*e*.*g*., *Hsp70Aa*), were excluded from our analysis. Among the 86 genes in the Heat Shock Proteins group (HSP; FBgg0000501), we detected 67 total HSP transcripts. Maternally deposited HSP transcripts (n = 62) were not enriched in the selection hits compared to the background set of genes (Fisher’s exact test, p = 0.64). However, zygotically transcribed HSPs (n = 64) were enriched in the selection hits compared to the background set (Fisher’s exact test, p = 0.0062; **Fig. 4A**), with 7 directionally selected HSPs (7 / 368, 1.9%; Fisher’s exact test, odds ratio = 2.65 [95% CI 1.01–5.90], p = 0.027), while stabilizing selection hits were not statistically enriched (2 / 60, 3.3%; Fisher’s exact test, odds ratio = 4.71 [95% CI 0.54–18.6], p = 0.074). It is interesting to note that parental bias in directional selection appeared to be stage-specific with 5 out of the 6 directionally selected maternal HSPs with higher expression in St. Kitts (**Fig. 4B**), and 6 out of 7 directionally selected zygotic HSPs with higher expression in Vermont (**Fig. 4C**). These directionally selected HSPs have a variety of roles in early embryos; *ClpX* and *Gp93* are involved in embryonic gut development, while small heat shock protein 23 (*Hsp23*) and the HSP70 superfamily genes (*Hsp68, Hsp70Ab*, and *Hsp70Bbb*) are known to be responsive to heat stress.

**Figure 4.**
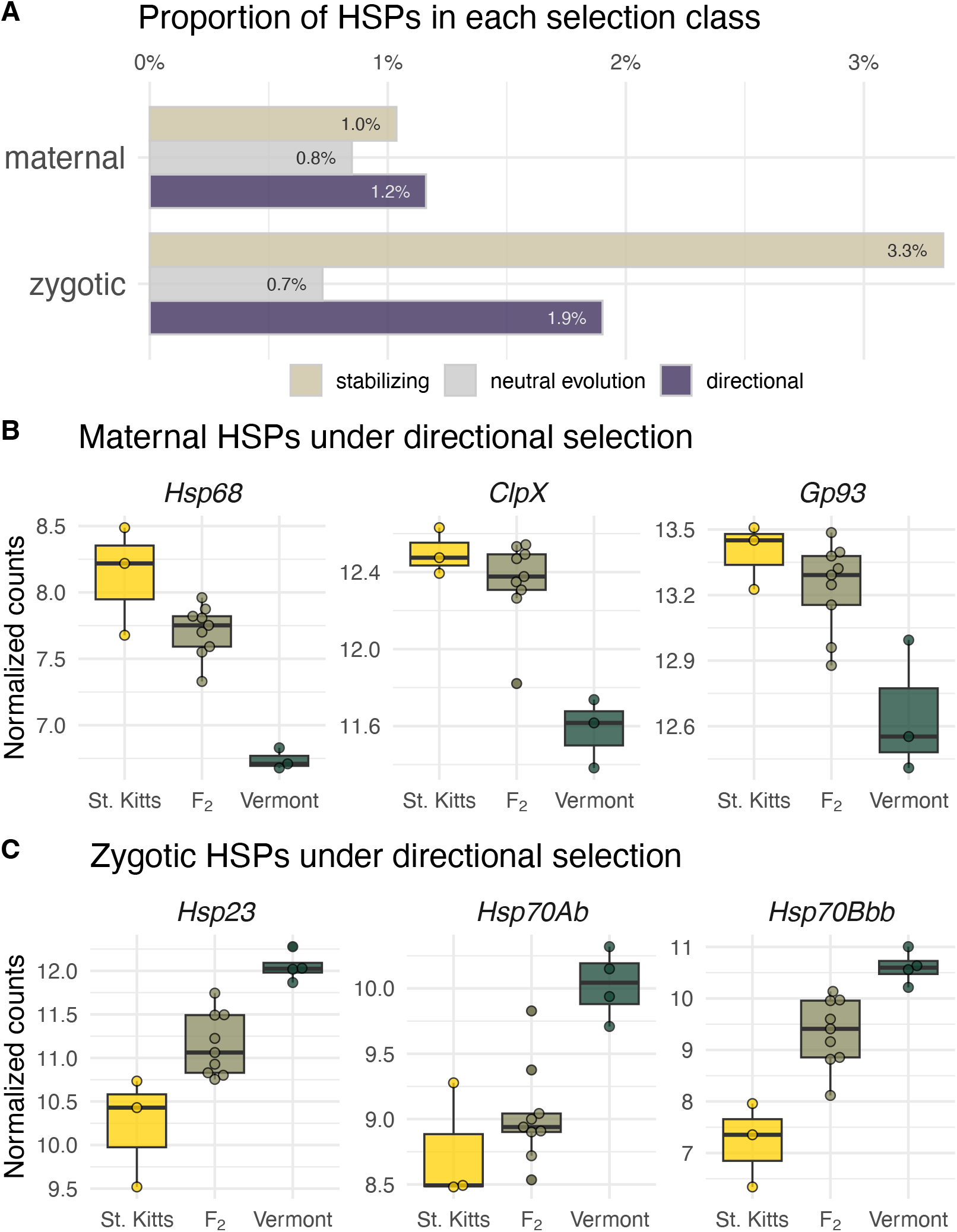
Heat-shock proteins show stage-specific patterns: maternal HSPs skew towards higher expression in St. Kitts, and zygotic HSPs skew towards increased expression in Vermont and are enriched among the selection hits. (**A**) Proportion of HSPs among maternal and zygotic genes in each selection class. (**B**) Maternal expression of three HSPs (*ClpX, Gp93*, and *Hsp68*) directionally selected for higher expression in St. Kitts. (**C**) Zygotic expression of three HSPs (*Hsp23, Hsp70Ab*, and *Hsp70Bbb*) genes directionally selected for increased expression in Vermont embryos. Normalized expression values for individual embryos (points) overlaid on top of box plots (boxes represent the interquartile range; middle bar indicates the median; whiskers extend to 1.5×IQR).

### Non-additive genetic effects

Finally, we tested for non-additive genetic effects on all genes (Fraser, 2020). Of 710 maternally expressed genes under selection, 91 (12.8%) showed significant non-additivity, with an additional 432 (6.9% of the 6,242) neutrally evolving genes showing non-additive genetic effects (adj. p < 0.05; **Fig. S9A**). No zygotic genes exhibited non-additivity (**Fig. S9B**). Non-additive genetic effects observed in the F_2_ segregants were not distributed proportionately across selection classes (Fisher’s exact test, p = 5.4 × 10^−9^); instead, directionally selected genes were roughly twice as likely to show non-additive genetic effects (14.7%; 76 / 517) as neutrally evolving genes (6.9%; 432 / 6242; Fisher’s exact test, odds ratio = 2.32 [95% CI 1.76–3.02], p = 5.4 × 10^−9^), whereas genes under stabilizing selection had a similar rate of non-additive genetic effects (7.8%, 15 / 193; Fisher’s exact test, odds ratio = 1.13 [95% CI 0.616–1.94], p = 0.67; **Fig. S10A**). Non-additive F_2_ segregant means were just as likely to be above or below the expected-under-additivity midparent mean, indicating that non-additivity did not systematically bias expression upward or downward (**Fig. S10B**). However, non-additive F_2_ segregant means were biased towards the St. Kitts parental mean: 88.1% of directionally selected genes (67 / 76) were biased towards St. Kitts versus 11.8% (9 / 76) towards Vermont (**Fig. 5A**). The gene expression of *Cyp6a22, gd, Bicra*, and *Nft-2* illustrate this parental bias in F_2_ means (**Fig. 5B**).

**Figure 5.**
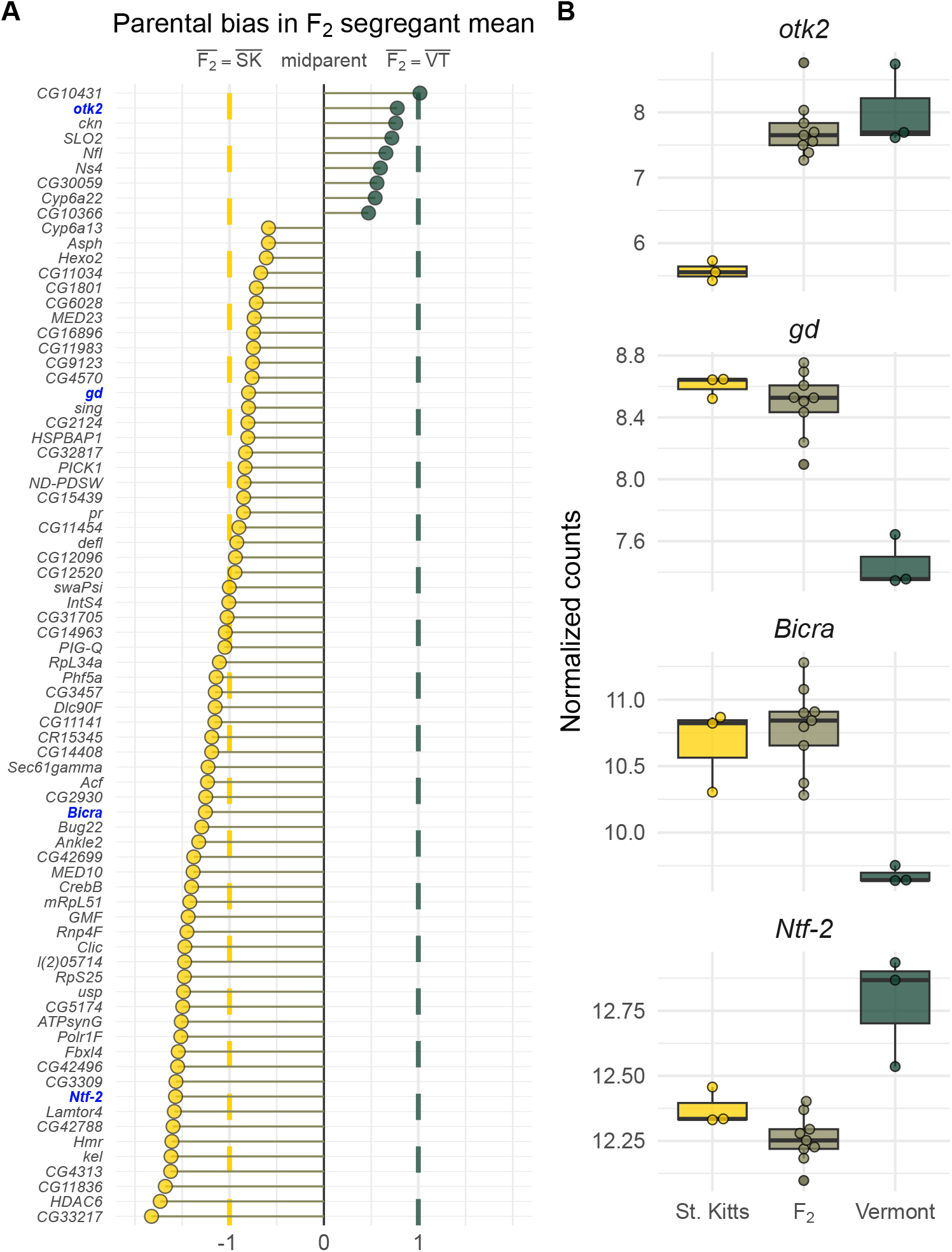
Among directionally selected maternal genes with non-additive inheritance, F_2_ means overwhelmingly bias towards the St. Kitts parental mean. (**A**) Dot plot of the parental bias index (PBI) for directionally selected maternal genes with non-additivity, organized by descending PBI. Positive values indicate the F_2_ mean is biased towards Vermont, and negative values indicate bias towards St. Kitts. Points are colored by parental bias of F_2_. The vertical dashed lines are where the F_2_ segregant mean is equal to the parental mean (PBI = -1 or 1). Values more extreme than these lines indicate cases where the F_2_ mean is outside of the range of the parental means. Gene symbols in blue indicate genes highlighted in the right panel. (**B**) Normalized expression of four non-additive genes, ordered by bias from Vermont to St. Kitts. Normalized expression values for individual embryos (points) overlaid on top of box plots (boxes represent the interquartile range; middle bar indicates the median; whiskers extend to 1.5×IQR).

## Supplementary Figures

**Figure S1.**
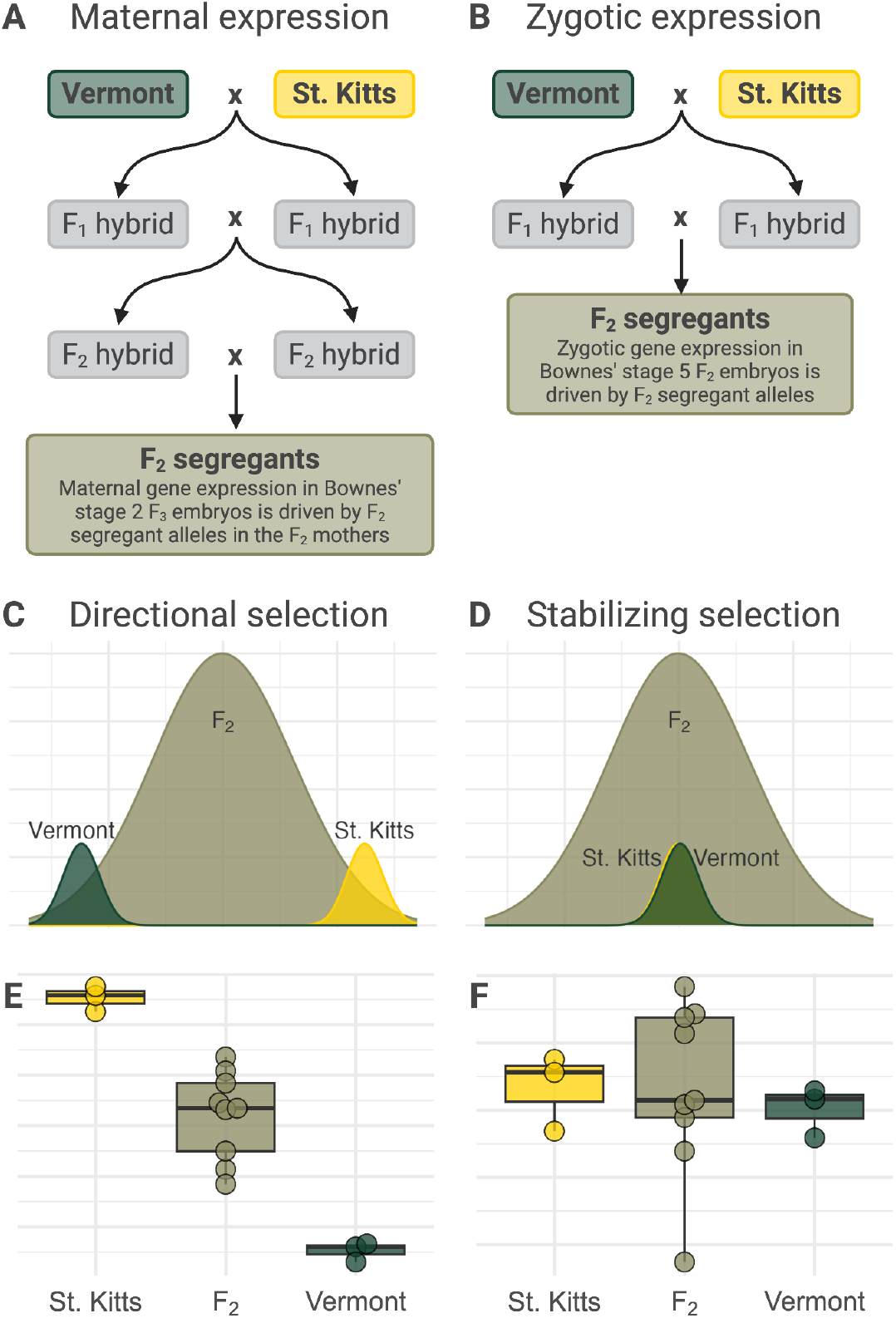
Genetic crossing design and visual representations of the variance test. Schematic of the genetic crossing designs used to generate F_2_ segregants to detect selection on maternal gene expression in Bownes’ stage 2 embryos (**A**), and zygotic expression in Bownes’ stage 5 embryos (**B**) using the variance test. Theoretical population distributions showing directional selection (**C**), and stabilizing selection (**D**), with a boxplot showing hypothetical data sampled from those distributions under directional (**E**) or stabilizing selection (**F**).

**Figure S2.**
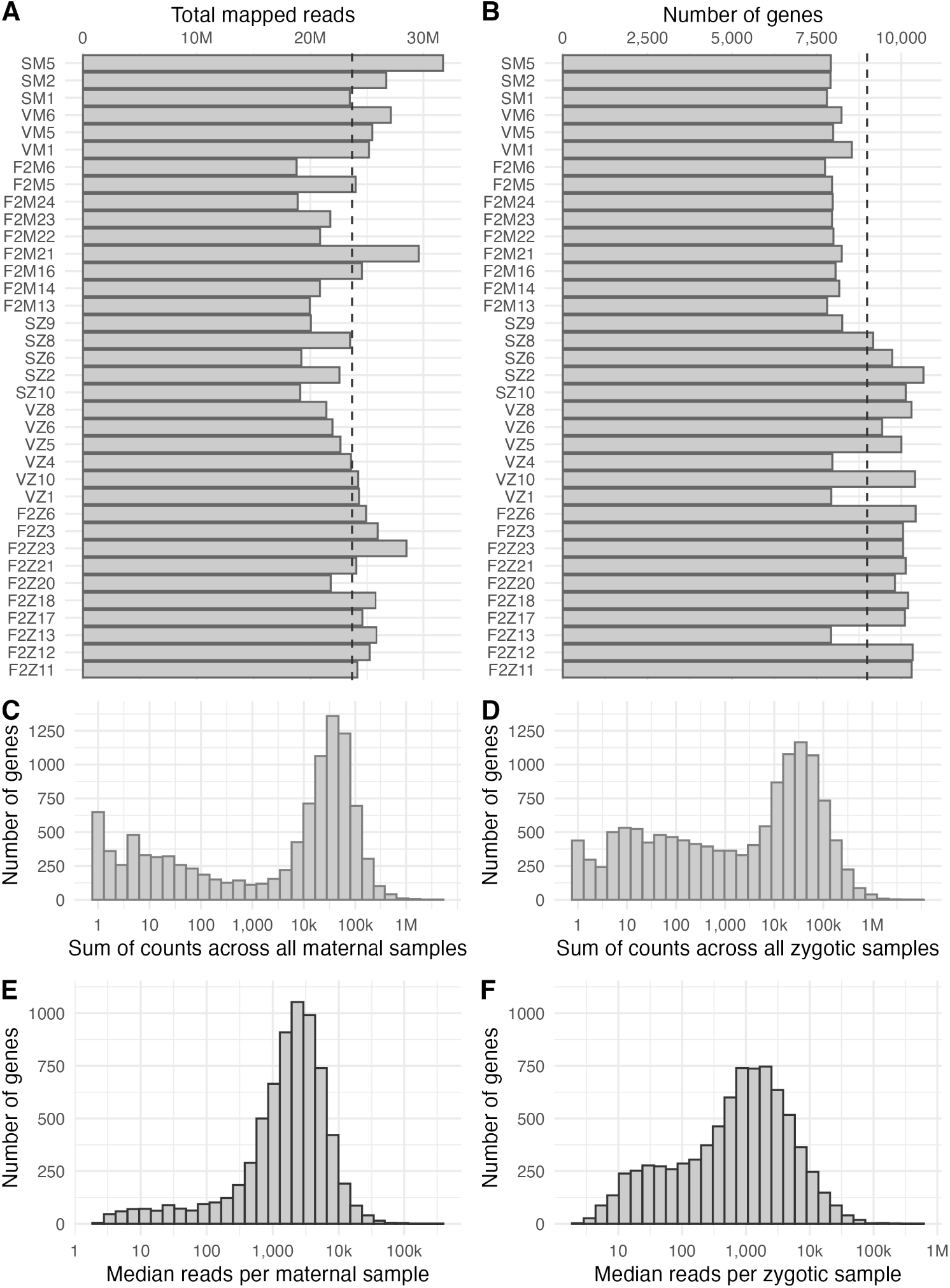
Sequencing quality control metrics. (**A**) Total number of mapped reads (millions) per sample, with mean (vertical dashed line). (**B**) Total number of genes detected per sample, with mean (vertical dashed line). (**C–D**) A histogram of per gene total number of counts across all maternal (**C**) and zygotic (**D**) samples. (**E–F**) A histogram of median reads per gene (included in variance tests) for maternal (**E**) and zygotic (**F**) samples.

**Figure S3.**
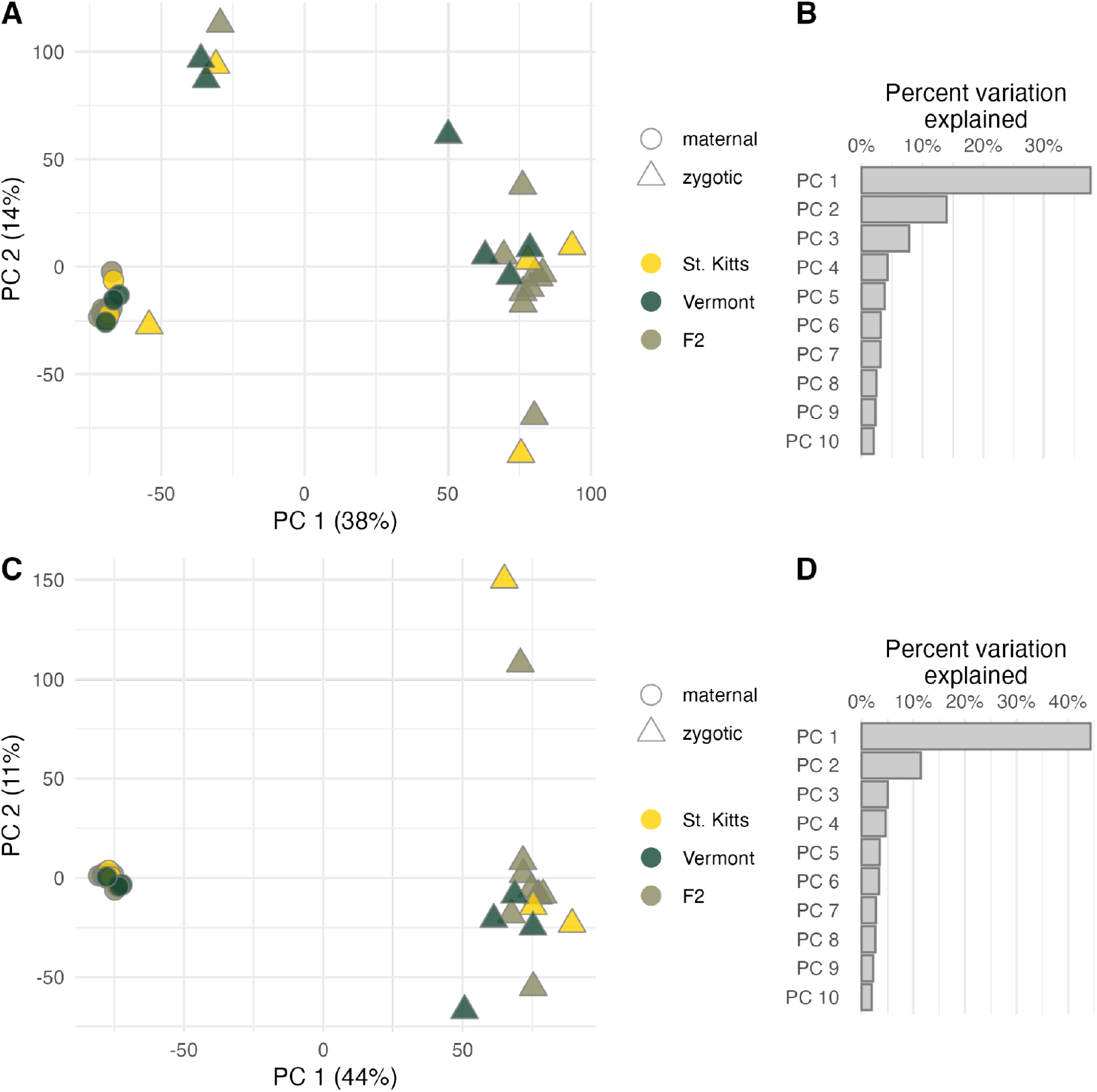
PCA for all samples with and without the anomalous samples. Principal component analysis of all 36 samples (**A**) and the corresponding scree plot showing the percent variation explained by the first 10 principal components (**B**). Principal component analysis of after removing the five anomalous zygotic samples (**C**) and the corresponding scree plot showing the percent variation explained by the first 10 principal components (**D**).

**Figure S4.**
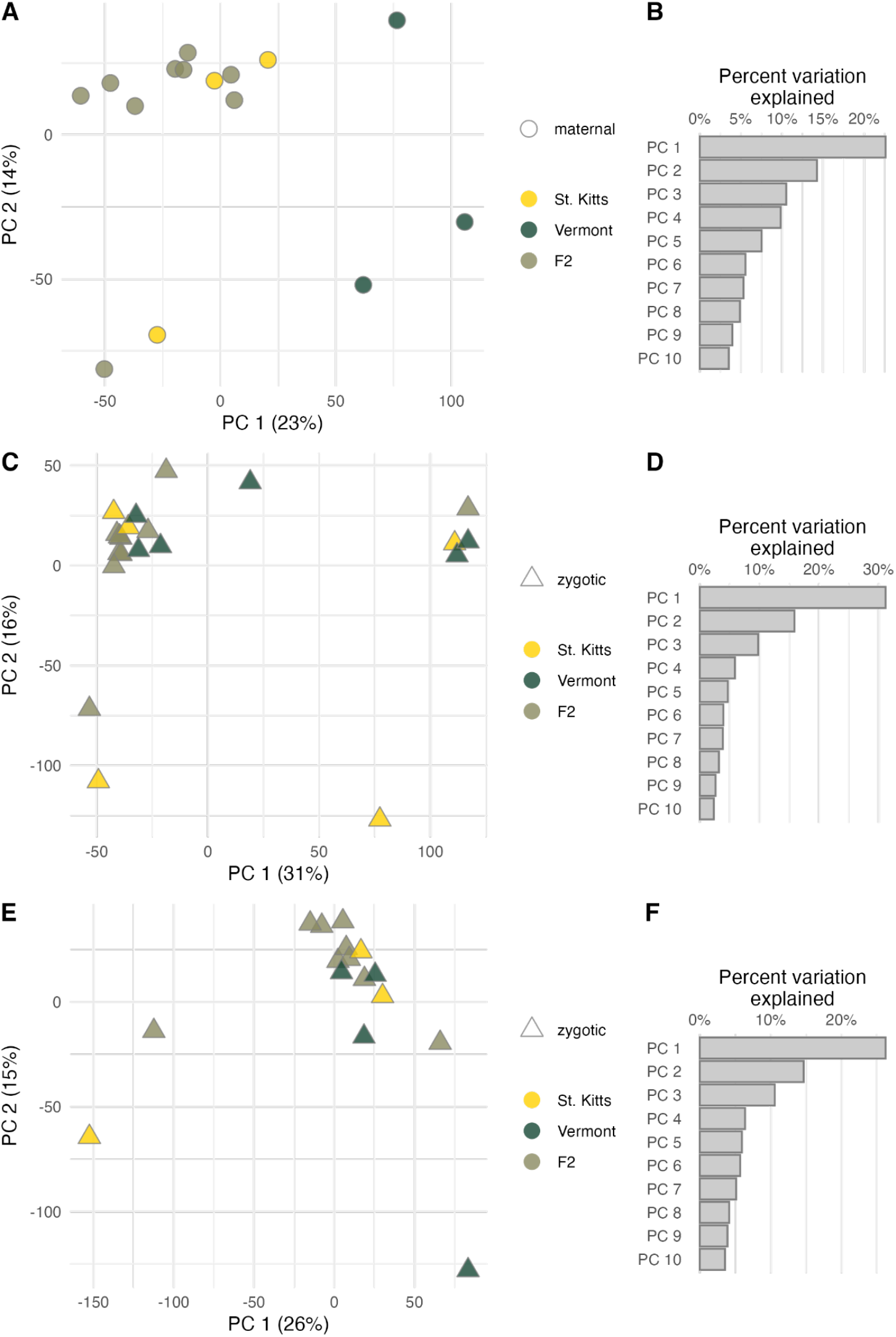
Stage-specific PCA for all samples with and without the anomalous samples. Principal component analysis of all 15 maternal samples (**A**) and the corresponding scree plot showing the percent variation explained by the first 10 principal components (**B**). Principal component analysis of all 21 zygotic samples (**C**) and the corresponding scree plot showing the percent variation explained by the first 10 principal components (**D**). Principal component analysis of the zygotic samples after removing the five anomalous samples (**E**) and the corresponding scree plot showing the percent variation explained by the first 10 principal components (**F**).

**Figure S5.**
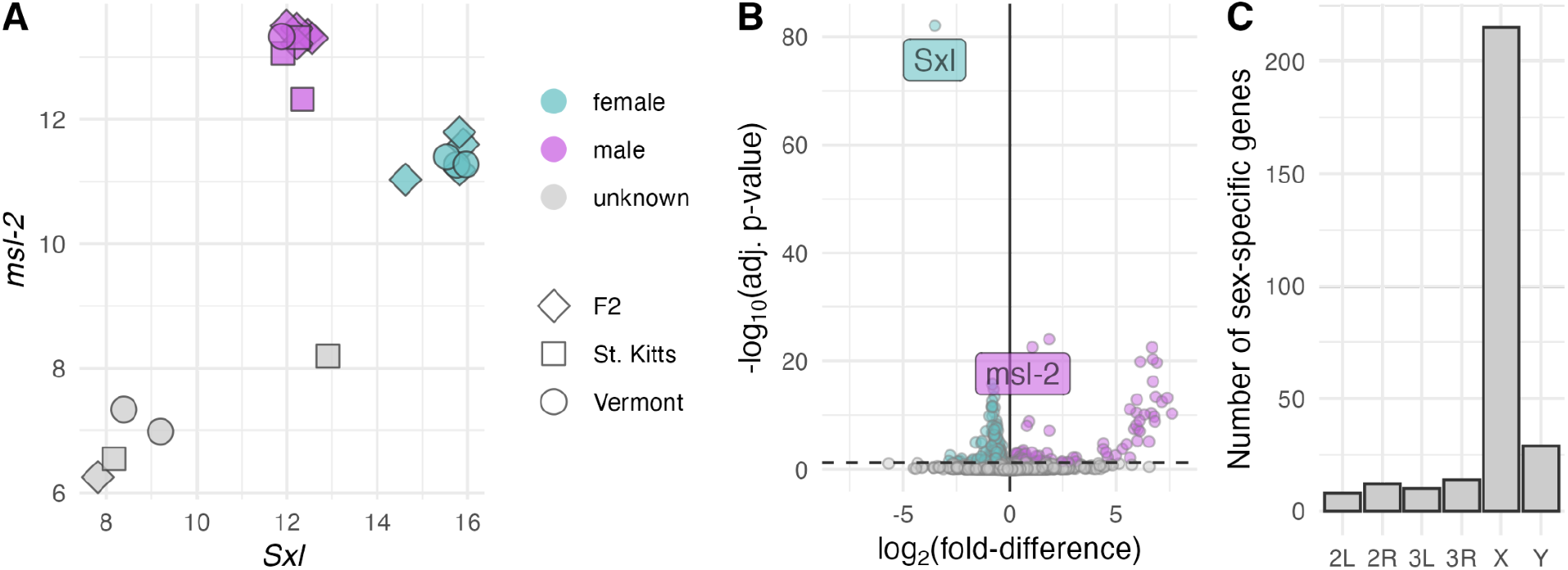
Sex-specific expression of Bownes’ stage 5 embryos. (**A**) Scatter plot of normalized *Sxl* (horizontal axis) versus *msl-2* (vertical axis) expression. Colored by assigned sex; low-expressing samples are labeled unknown (five anomalous samples). (**B**) DESeq2 volcano log_2_(fold-difference) versus -log_10_(adj. p-value). Sex-specific DEGs (adj. p < 0.05) are colored by sex bias (male-biased genes in lavender and female-biased genes in teal). (**C**) Bar plot sex-specific gene counts per chromosome.

**Figure S6.**
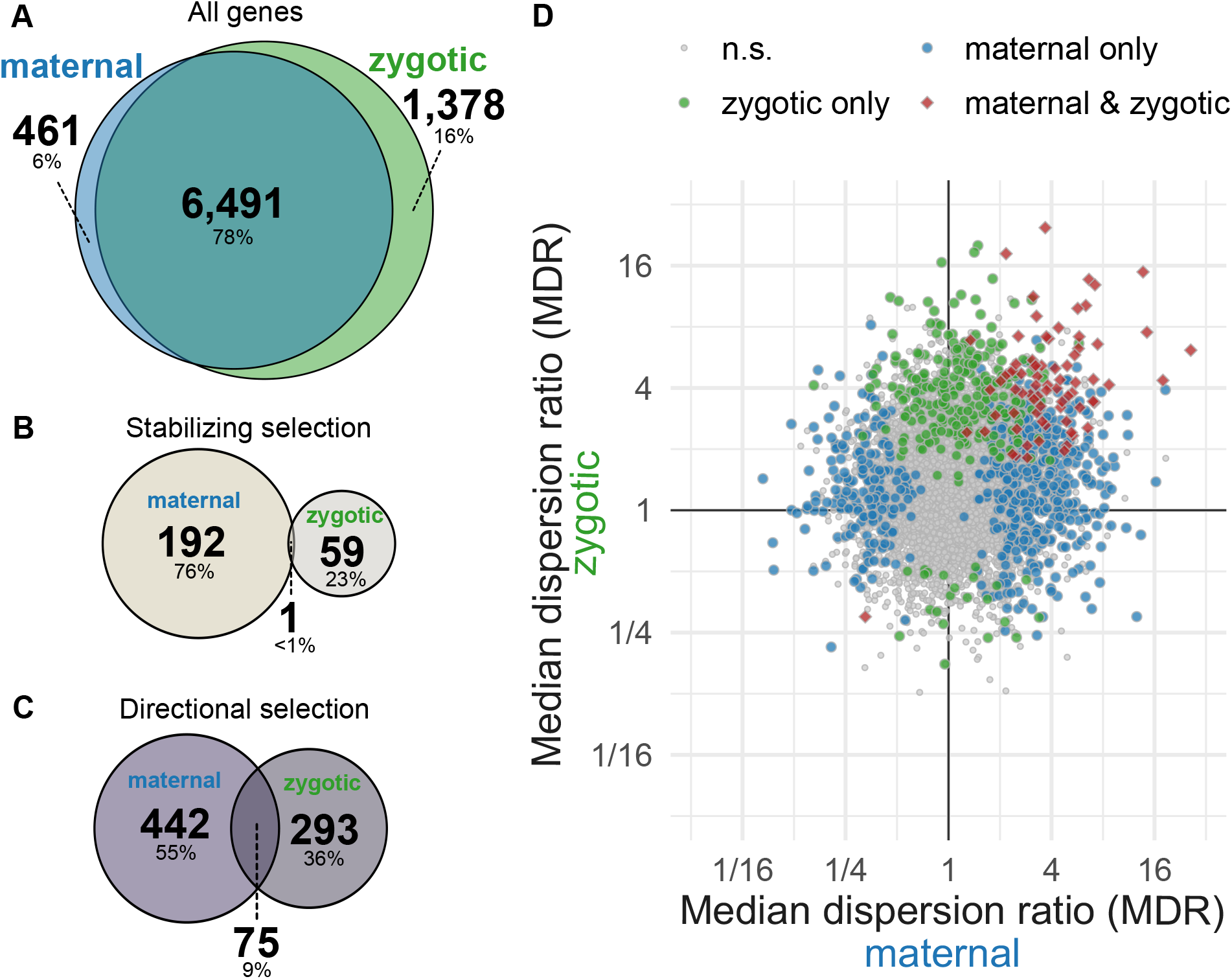
Shared and stage-specific expression and selection. (**A**) Venn diagram of all genes tested for selection in maternal (Bownes’ stage 2) and zygotic (Bownes’ stage 5) expression. (**B**) Venn diagram of genes with significant stabilizing selection in maternal and zygotic stages. (**C**) Venn diagram of genes with significant directional selection in maternal and zygotic stages. (**D**) Scatter plot of MDR in maternal (horizontal axis) versus zygotic expression (vertical axis). Points are colored by selection significance: neutral at both stages (grey), maternal only (blue), zygotic only (green), and both stages (red).

**Figure S7.**
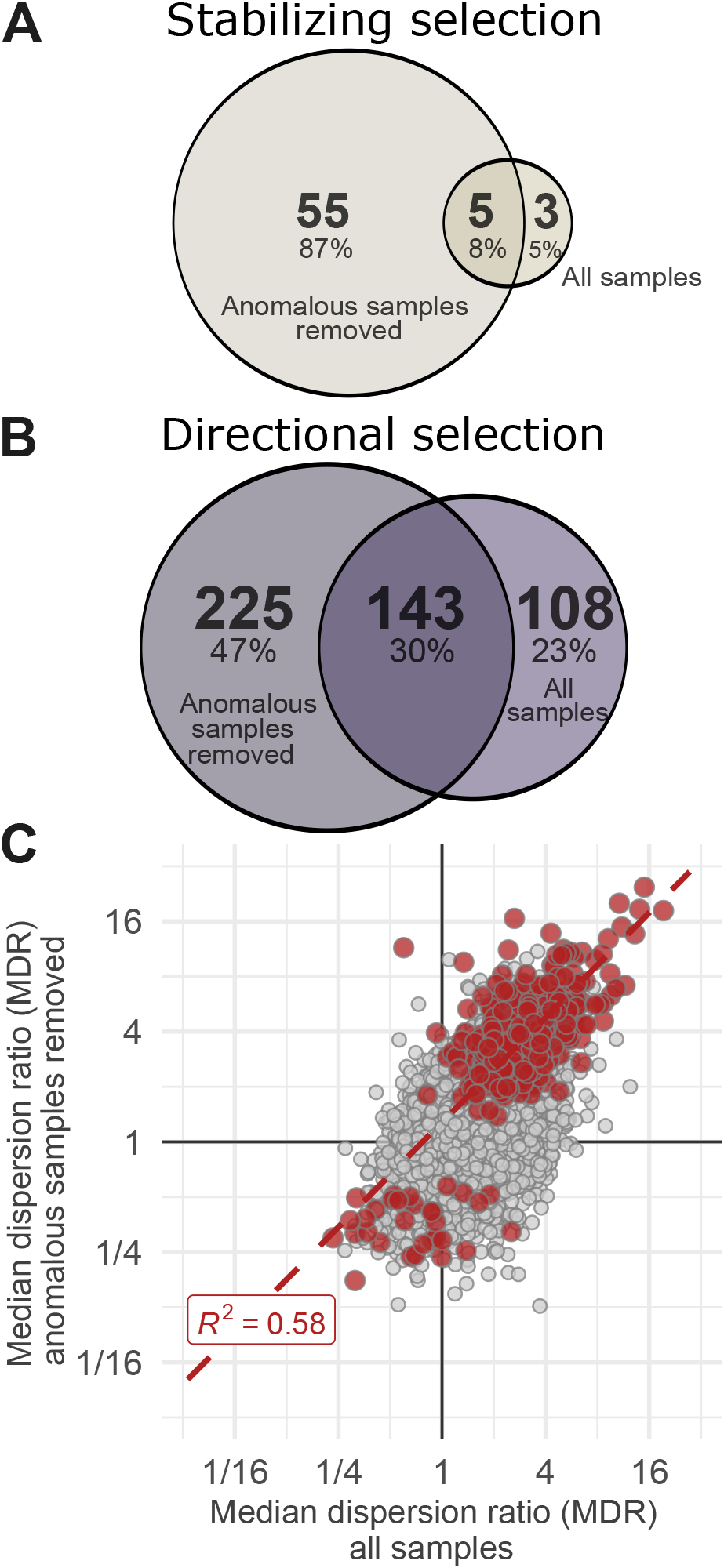
Detecting selection on zygotic expression levels with and without the five anomalous samples. (**A**) Venn diagram of stabilizing-selection genes identified with and without the five anomalous samples. (**B**) Venn diagram of directional selection genes identified with and without the five anomalous samples. (**C**) Scatter plot comparing median dispersion ratios (MDR) for each gene between the two analyses; red points indicate genes under significant stabilizing or directional selection (in the filtered analysis), and the dashed line shows the linear fit of the genes under selection and corresponding R^2^.

**Figure S8.**
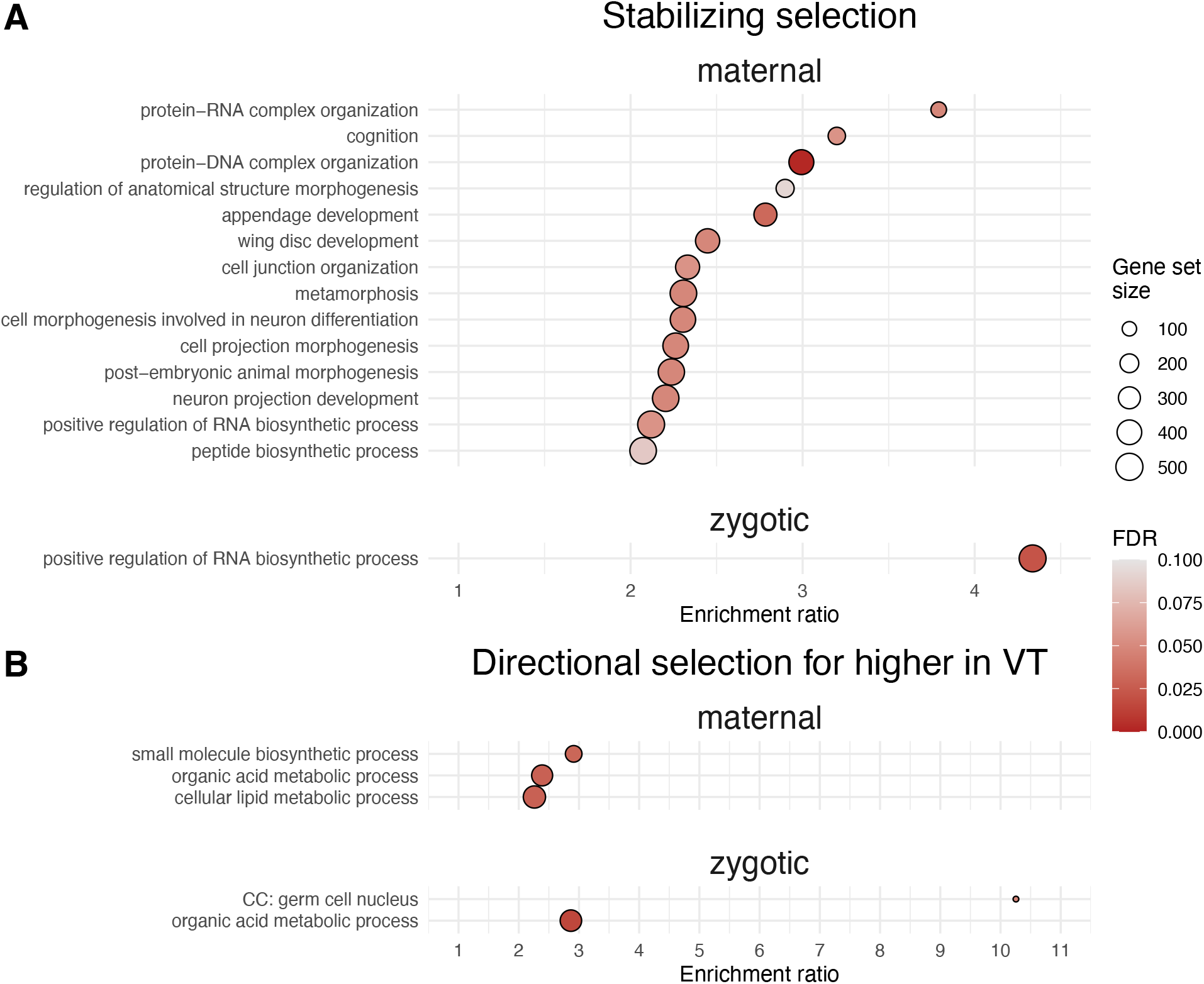
Over-representation analysis of gene ontology (GO) terms in the selection results. (**A**) GO terms over-represented in genes under stabilizing selection at maternal and zygotic stages. (**B**) GO terms over-represented in genes directionally selected for higher expression in Vermont at maternal and zygotic stages. (No GO terms were over-represented in genes with directional selection towards increased expression in St. Kitts). Point size denotes the number of genes in each GO set and color indicates FDR. The horizontal axis represents the enrichment ratio (*i*.*e*., observed vs. expected DEGs). All terms with FDR < 0.1 are shown.

**Figure S9.**
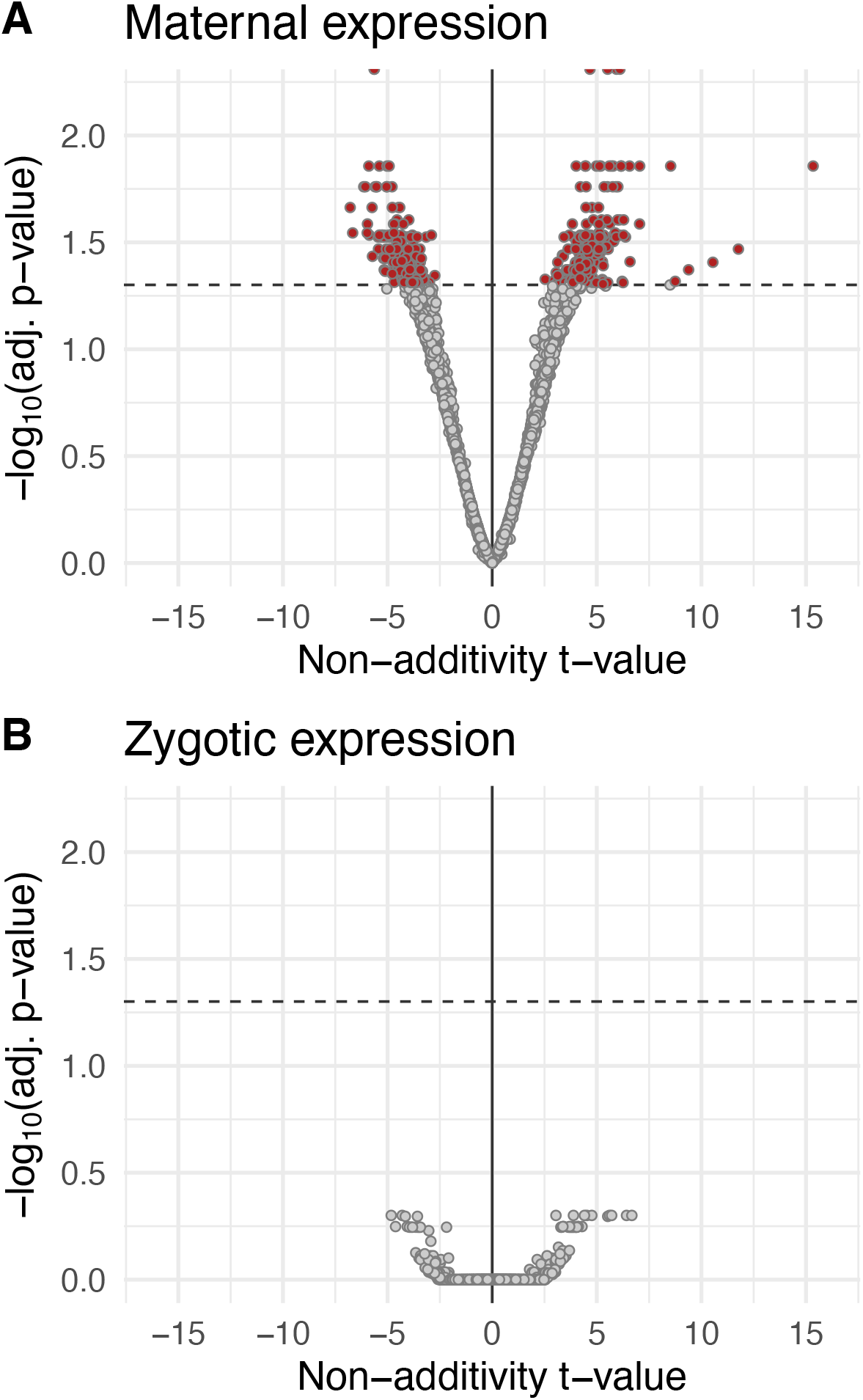
Non-additive genetic effects in maternal and zygotic expression. Volcano plot of test for genetic non-additivity in the maternal (**A**) and zygotic (**B**) genes. Horizontal axis shows the direction and magnitude of non-additive effects (positive if F_2_ mean > midparent expectation; negative if F_2_ mean < midparent expectation), and the vertical axis indicates the significance (-log_10_(adj. p-value)). Red points indicate significant non-additive genetic effects (adj. p < 0.05).

**Figure S10.**
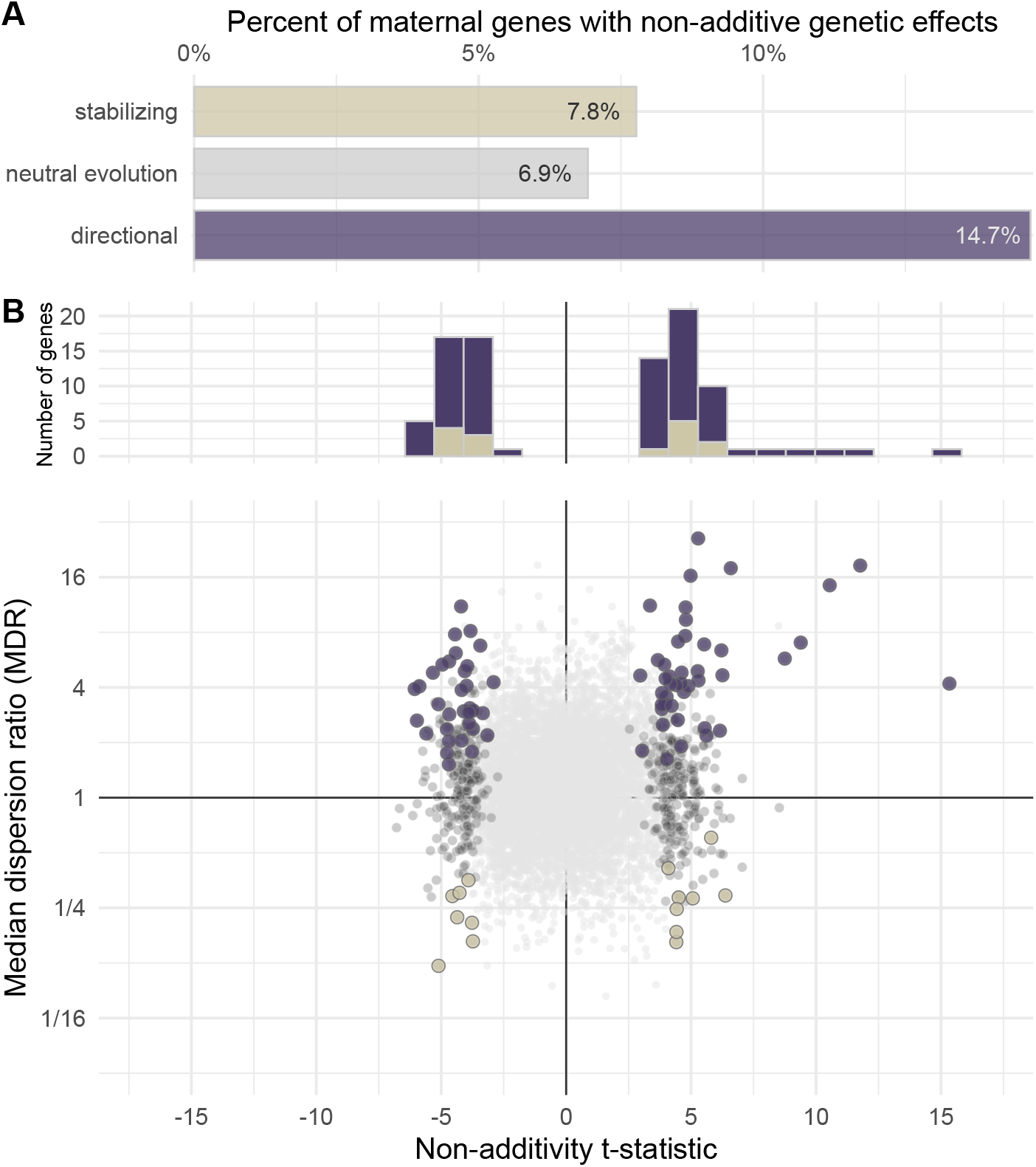
Non-additive genetic effects in the maternally expressed genes. (**A**) Bar plot showing the percentage of maternal genes with non-additive genetic effects in each selection class. (**B**) Top: histogram of non-additivity t-statistic for maternally selected genes with non-additivity. Bottom: scatter plot of median dispersion ratio (MDR) versus non-additivity t-statistic. Points are colored by selection class and non-additivity: purple for directional selection genes with non-additive genetic effects, tan for stabilizing selection genes with non-additivity, dark gray for non-additive genes under neutral evolution, and light gray for genes with neither non-additivity nor selection.

## Discussion

Using genetic crosses between tropical (St. Kitts) and temperate (Vermont) lines, we analyzed F_2_ segregants with a cross-based variance test (Fraser, 2020) to infer selection on gene expression at maternal (Bownes’ stage 2) and zygotic (Bownes’ stage 5) developmental timepoints. The tension between the highly conserved maternal-to-zygotic transition (MZT) (Tadros and Lipshitz, 2009; Vastenhouw et al., 2019) and the status of early embryos as a target of thermal selection (Lockwood et al., 2018) makes this developmental window an especially compelling context to examine selection on gene expression. We find stage-specific signals of selection on expression, with directional selection more common, especially among maternally deposited genes, and a high degree of stabilizing selection on core zygotic regulators.

Contrary to our expectations, signals of directional selection were more prevalent than stabilizing selection, with about twice as many directional selection genes as stabilizing selection genes in the maternal transcriptome and approximately five times as many in the zygotic stage. This result conflicts with the canonical view from comparative, mutational, and modeling studies that selection on gene expression is dominated by stabilizing forces, constraining natural variation in the transcriptome (Bedford and Hartl, 2009; Gilad et al., 2006; Rifkin et al., 2003; Rifkin et al., 2005; Romero et al., 2012). This apparent discrepancy may be explained by disparate evolutionary time scales between experiments (Price et al., 2022). Comparative work relies on inter-specific differences accumulated over millions of years of evolution, whereas our study captures ongoing within-species dynamics (Fraser, 2020). It is plausible that stabilizing selection dominates over longer evolutionary periods, as the microevolutionary timescale we sampled here does not capture the historical canalization events that have already been fixed in the distant past (Flatt, 2005; Lott et al., 2007; Siegal and Bergman, 2002; Waddington, 1942; Waddington, 1959). Furthermore, given evidence that selection has shaped thermal tolerance in heat-sensitive early embryos (Lockwood et al., 2018), an excess of directional selection may be consistent with local adaptation in gene expression. Overall, the 5–10% of genes under selection we report likely represents a conservative lower bound and is not directly comparable to the proportions reported in inter-specific divergence studies.

As predicted, maternally deposited transcripts showed more widespread signals of selection than zygotic transcripts; specifically, we detected signals of selection in 10.2% of maternal-stage transcripts compared with only 5.4% of zygotic-stage transcripts. This recapitulates the hourglass model of developmental divergence (Duboule, 1994; Von Baer, 1828) and aligns with reported patterns of embryonic gene expression across a diverse array of taxa (Atallah and Lott, 2018; Irie and Kuratani, 2011; Kalinka et al., 2010; Quint et al., 2012). Much of this previous work did not distinguish adaptive divergence from neutral drift. More recently, molecular evolution analyses have shown that maternal genes in *Drosophila* accumulate higher levels of non-adaptive, neutral divergence than zygotic genes in protein-coding regions (Coronado-Zamora et al., 2019), but that levels of adaptive evolution are similar. This suggests that maternal genes are less evolutionarily constrained than zygotic genes but not specifically targeted by directional or stabilizing selection within their protein-coding regions. However, our expression data demonstrate clear signals of directional selection and stabilizing selection at the transcript level in maternal genes. Thus, while coding sequence variation accumulates primarily via neutral drift in maternal genes, selection fine-tunes their regulatory variation.

Interestingly, within the zygotic transcriptome, zygotic-only genes (*i*.*e*., genes absent at the maternal stage) were disproportionately represented among the stabilizing selection hits, suggesting that genes activated during zygotic genome activation are particularly likely to be under stabilizing selection. These stage-specific patterns of selection are broadly consistent with prior comparative work across the *Drosophila* phylogeny, which has shown the greatest conservation in the expression of zygotic-only genes (Atallah and Lott, 2018; Feitzinger et al., 2022).

The regulatory targets of selection are likely stage-specific. Previous work shows that interspecific differences in maternal transcription are primarily driven by *trans*-regulatory mutations, whereas zygotic differences arise from a combination of *cis* and *trans* mutations (Cartwright and Lott, 2020). This genetic architecture mirrors the underlying regulatory environments of the two stages: the maternal genome is largely under the control of *trans*-acting modifiers of chromatin state, while the zygotic genome is regulated by gene-specific transcription factors (Omura and Lott, 2020). Because the transcripts of these modifiers of chromatin state are largely present in maternal nurse cells supporting oogenesis, with only the protein loaded into the embryo (Berg et al., 2024; Crofton et al., 2018; Mische et al., 2007), their expression is not directly visible to our test of selection. This stage-specific difference in the relative magnitude of *cis* versus *trans* gene regulation is important to note in the context of our cross-based test of selection, which relies on random recombination of parental alleles in the F_2_ segregants. Because *trans*-acting alleles are often unlinked to their various targets, recombination shuffles them into novel allele combinations, generating new additive and non-additive (epistatic) *trans*-*trans* combinations that may broaden the expression variation among many target genes, and inflate stabilizing signals across the transcriptome. In contrast, *cis*-acting alleles remain anchored to their target genes after recombination and are therefore less likely to alter the transcriptome-wide F_2_ distribution. On the other hand, directional selection on a small number of *trans*-acting maternal regulators could drive a disproportionately large number of directional selection hits in the maternal transcriptome simply due to network amplification on the expression of downstream effectors. The balance of these effects depends on the relative effect sizes of *cis* versus *trans* alleles on expression differences.

Selection on the expression of transcription factors (TFs) and canonical embryogenesis genes showed stage-specific patterns. Canonical zygotically transcribed genes and TFs were enriched for stabilizing selection and under-represented in directional selection hits relative to their representation among neutrally evolving genes. This zygotic result aligns with extensive research showing that zygotic regulators are highly canalized through enhancer redundancy (buffering against mutational perturbations) and the cross regulation of gap genes (Atallah and Lott, 2018; Gaskill et al., 2021; Manu et al., 2009; Perry et al., 2010). For example, short gastrulation (*sog*), a bone morphogenic protein (BMP) pathway inhibitor involved in enforcing the Dpp gradient central to dorsal-ventral patterning (Eldar et al., 2002), is one of many zygotic genes with shadow enhancers that support developmental robustness (Chopra et al., 2012; Perry et al., 2010; Shin and Hong, 2016). Additionally, among the stabilizing-selection genes are two anterior-posterior regulators caudal (*cad*) and ocelliless (*oc*), repressed and activated, respectively, by maternal Bicoid protein. These opposing expression gradients, together with other gap genes (*e*.*g*., hunchback, Kruppel, giant, and knirps) create cross-repression that enforces tight canalization of gene expression during zygotic genome activation (Finkelstein and Perrimon, 1990; Jaeger, 2011; Rivera-Pomar et al., 1996). Although, we do detect directional selection on some zygotic regulators (*e*.*g*., *Paris* and *fuss*), they are under-represented among the directional hits.

Contrary to expectations, canonical maternal genes and TFs showed no enrichment among our selection hits. One plausible explanation is that long-term stabilizing selection has limited the standing genetic variation such that insufficient variation exists between the genotypes we crossed to reveal new stabilizing selection at these core maternal genes (Bedford and Hartl, 2009). Alternatively, the lack of selection enrichment in core maternal genes may be due to these genes being primarily regulated post-transcriptionally (Benoit et al., 2009; Wang et al., 2017). For example, key axis-patterning maternal RNAs oskar and nanos are kept translationally silent by ribonucleoprotein (RNP) repressors and RNA-binding proteins (*e*.*g*., Smaug), and translation is possible only after correct localization (Chen et al., 2014; Kim-Ha et al., 1995; Smibert et al., 1996), suggesting transcript abundance is often not the limiting factor. Finally, our whole embryo bulk RNA data does not capture localization of transcripts, so any spatial co-localization involved in canalization is invisible to our test of selection (Ding et al., 1993; Kugler and Lasko, 2009; Lasko, 2012).

Signals of local adaptation in HSP expression were stage-dependent, with opposite directions of selection in maternal- and zygotic-stage embryos. In heat-tolerant embryos from St. Kitts, we observed directional selection towards higher maternal loading of HSPs, including *Hsp68* (an inducible Hsp70-family chaperone), *ClpX* (a mitochondrial chaperone), and *Gp93* (an Hsp90-family chaperone involved in gut development) (Li et al., 2025; Matsushima et al., 2017; Maynard et al., 2010). This matches prior work showing that increased maternal deposition of molecular chaperones can increase early embryonic heat tolerance (Lockwood et al., 2017), which is especially relevant given the lack of zygotic transcription prior to ZGA (Vastenhouw et al., 2019). This finding is in contrast with previous work which failed to find differences in the expression of HSPs between heat-sensitive and heat-tolerant genotypes (Mikucki et al., 2024), although these embryos were assayed later in development and had a mix of maternal and zygotic transcripts. In contrast, the zygotic stage embryos revealed directional selection for increased HSP expression (*e*.*g*., *Hsp23, Hsp70Ab*, and *Hsp70Bbb*) in Vermont, a pattern that seems counter to their observed heat-sensitivity. However, Bownes’ stage 5 occurs a couple of hours after the most heat-sensitive stage in embryonic development, and regional differences in heat tolerance disappear later in development (Lockwood et al., 2018). Finally, we note that our baseline (25°C) expression measurements do not capture any differences between genotypes in the thermal responsiveness of HSPs.

We also detected directional selection for higher Vermont expression of genes annotated to the germ cell nucleus. These genes, including *piwi, rhi*, and *vas*, are involved with the PIWI-interacting RNA (piRNA) pathway, which suppresses transposon activity, protecting germline cells from DNA damage (Czech et al., 2018; Huang et al., 2017; Ozata et al., 2019), and together with Hsp90 they have been hypothesized to maintain developmental robustness (Rohner et al., 2013; Rutherford and Lindquist, 1998; Specchia et al., 2010). Deposited maternally, these genes are also transcribed zygotically to buffer against increased transposable element (TE) transcription which peaks several hours after fertilization (Fabry et al., 2021). While there is no evidence of latitudinal differentiation in TE abundance or allele frequencies in *D. melanogaster* (Adrion et al., 2019), because this is a temperature-sensitive system (Ho et al., 2024), higher expression of piRNA pathway genes in Vermont may buffer against the high thermal variation experienced by temperate embryos (Hoffmann et al., 2003; Roberts and Feder, 2000).

Finally, we did not observe any significant non-additive genetic inheritance in the zygotic-stage F_2_ segregants. The absence of non-additivity again supports the idea that zygotic transcription, with its cross-regulatory dynamics and redundant enhancers is a highly canalized, robust developmental system (Jaeger, 2011; Manu et al., 2009; Perry et al., 2010). In contrast, the maternal F_2_ segregants showed hundreds of genes with non-additive inheritance; and among those directionally selected genes, the vast majority of F_2_ means were skewed towards St. Kitts. These non-additive effects are likely the consequence of epistasis (*i*.*e*., gene × gene interactions), dominance (*i*.*e*., the presence of one variant masking another) or both (Bourg et al., 2024; Gilchrist and Nijhout, 2001; McManus et al., 2010; Phillips, 2008). Because the maternal transcriptome is predominately regulated by chromatin state and other *trans*-acting effects (Cartwright and Lott, 2020; Omura and Lott, 2020), the skew towards St. Kitts could plausibly reflect the impact of a small set of *trans*-acting factors. This bias towards tropical parental expression mirrors the tropical bias in acute heat tolerance observed in F_1_ reciprocal crosses (Lockwood et al., 2018). In both cross directions, F_1_ embryos matched their tropical parent’s heat tolerance, indicating directional dominance of tropical alleles underlying this trait. However, because LT_50_ did not differ between reciprocal crosses, the mechanism underlying heat tolerance cannot be solely maternal, and must require zygotic input.

Together, our findings show that selection shapes embryonic transcript levels over short evolutionary timescales, but with different effects on the maternal and zygotic transcriptomes. As predicted, maternal transcripts showed more widespread signals of selection overall, but contrary to our expectations, genes with signals of directional selection outnumbered genes with signals of stabilizing selection. The enrichment of stabilizing selection in core zygotic regulators and the prevalence of directional selection among maternal molecular chaperones and germline defense genes suggest that selection targets gene sets in stage-specific ways. These results underscore that gene expression is an active target of natural selection over timescales relevant to local adaptation.

## Data availability

All sequencing data is available on the Sequence Read Archive (PRJNA1462806) and all code at GitHub (https://github.com/tsoleary/mzt_selection).

## Acknowledgements

We would like to thank Colin D. Meiklejohn for helpful discussions.

## Funding

This work was funded by NSF grant IOS-1750322 to BLL.

